# Biophysical Basis of the *in vivo* ERG of the Mouse: Rod, RPE and Müller glial Cell Contributions To Dark and to Light-driven Changes in Sub-Retinal Space [K^+^]_o_

**DOI:** 10.1101/2025.11.09.687427

**Authors:** Gabriel Peinado Allina, Prescott Alexander, Kaitryn Ronning, Marie E. Burns, Edward N. Pugh

**Affiliations:** Center for Neuroscience; Department of Ophthalmology & Vision Science; Department of Cell Biology & Human Anatomy; Department of Physiology & Membrane Biology

**Keywords:** Kir4.1, Kir7.1, quinacrine, Vu590, extracellular K^+^

## Abstract

The subretinal space (SRS) is a contiguous extracellular volume bounded by the external limiting membrane (ELM) and the retinal pigment epithelium (RPE). Approximately 90% of mouse rod photoreceptor dark current circulates in the SRS, and rod photoresponses materially alter SRS concentrations of permeant ions, especially K^+^. Following intravitreal injection of drugs that block glutamatergic synapses, and drugs that inhibit Kir4.1 and Kir7.1 channels expressed in Müller and RPE cells respectively, we measured electroretinograms (ERGs) in response to light flashes that completely suppress rod dark current, and compared ERG *c*-waves with predictions of a biophysical model of extracellular current sources generated by rod, Müller and RPE cells. SRS K^+^ fluxes in the dark steady-state were dominated by rods – NKX (–6.6 mM s^-1^), K_v_2.1 channels (+5.4 mM s^-1^), NCKX (0.3 mM s^-1^) and HCN1 channel fluxes (1.0 mM s^-1^) – with much smaller contributions from Kir4.1 and Kir7.1 channels and trans-ELM and trans-RPE fluxes combining to zero net flux at resting SRS *K*_o_ of 5 mM (re 3.5 mM [K^+^] in blood plasma). In response to rod-saturating stimulation, *K*_o_ was predicted to decline to ∼3.5 mM with a half-time of ∼ 0.5 s, driven mainly by a hyperpolarization-dependent decline in K_v_2.1 current. The kinetics and amplitude of the *c*-wave are explained by the decline in SRS *K*_o_, which negatively shifts the Nernst potential of apical RPE Kir7.1 channels, increasing outward current that sinks through the paracellular resistance to basal Bestrophin and CFTR chloride channels causing a trans-epithelial potential of ∼ 2 mV.

## INTRODUCTION

The physiological function of electrically active cells *in vivo* is affected by their morphology, their ion channel expression patterns and the properties of the tissues in which they reside. Among the latter properties are the concentrations of permeant ions in the local extracellular space (ECS). Ion concentrations in the relatively large volume of blood plasma are homeostatically maintained by the kidney. However, the relatively small ECS of most neuronal tissue is separated from the blood by several barriers, including vascular endothelial cells, pericytes, and the endfeet of astrocytes. In the small ECS volumes of nervous system parenchyma, neuronal activity can materially alter ion activity, in particular that of K^+^, whose resting level is typically near that of blood plasma, 3.5 – 4.5 mM in mammals.

The inner and outer segments of the rod and cone photoreceptors are situated in the subretinal space (SRS) (Steinberg, 1985a; Steinberg, 1985b), and have a high resting membrane current density of ∼ 23 μA cm^-2^, which switches signs from outward in the inner segment to inward in the outer segment (Hagins et al., 1970; Peinado Allina, 2025b, Fig. S1). On the inner segment (proximal) side the SRS is bounded by the external limiting membrane (ELM), formed by Müller glial apical endfeet adjoined by adherens junctions (Daniele et al., 2007b). On its outer segment (distal) side the SRS is bounded by the retinal pigment epithelium (RPE), a monolayer of cells interlinked by tight and adherens junctions which form a major component of the blood-retinal barrier of the posterior eye (Jin et al., 2002). In darkness, a large inward Na^+^ current through CNG channels together with the inward electrogenic NCKX current is electrically balanced largely by efflux of K^+^ via electrogenic NKX transport and K_v_2.1 channels, both located exclusively in the inner segment (Fortenbach et al., 2021). Approximately 90% of the dark current of the densely packed photoreceptor population thus circulates within the restricted SRS, and consequently their light responses can cause functionally material changes in the extracellular concentrations of permeant cations, particularly K^+^. Specifically, investigations in amphibia and cat retina established that light stimulation of photoreceptors causes [K^+^]_o_ (hereafter, *K*_o_) in the SRS to decline from a relatively high resting level (∼ 5 mM) to 3 mM or less (Oakley and Green, 1976; Oakley, 1977; Steinberg et al., 1980; Steinberg, 1985b; Gallemore et al., 1997). A central hypothesis arising from these investigations, when combined with others of the electrical properties of the RPE (Adijanto et al., 2009), is that light-dependent decreases in SRS *K*_o_ are driven by net uptake of K^+^ by rod photoreceptors, and that the consequently diminished SRS *K*_o_ in turn causes a trans-epithelial potential observable in the corneal electroretinogram (ERG) as the *c*-wave. We identify this explanation of the *c*-wave as the Oakley-Green-Steinberg-Miller (OGSM) hypothesis, based on the original and extensive contributions of these investigators and their coworkers to the relevant physiology of the RPE and SRS.

The present investigation examines the OGSM hypothesis in the living mouse eye, using a gene-targeted mouse (*Gnat2*^−/−^ ) to eliminate cone responses (Ronning et al., 2018), and pharmacological manipulations to block glutamatergic synaptic transmission and to selectively inhibit K^+^ transport by Kir channels of the RPE (Kir7.1) and Müller cells (Kir4.1). The primary goal is to determine whether the OGSM hypothesis is quantitatively consistent with the molecular mechanisms established to underlie the dark state and photoresponses of mouse rod photoreceptors (Peinado Allina et al., 2017; Fortenbach et al., 2021; Peinado Allina, 2025b), and the wealth of evidence about the molecular mechanisms underlying the mammalian RPE’s electrical properties.

## MATERIALS and METHODS

### Animals and experimental protocol

#### Mice and husbandry

All mice were cared for and handled with approval of the University of California, Davis, Institutional Animal Care and Use Committee and in accordance with National Institutes of Health Guidelines. General methods used for the ERG recordings are provided in an accompanying papers (Peinado Allina, 2025a; Peinado Allina, 2025b), and here only methods specific to this investigation are presented.

#### Intravitreal injections

All procedures were performed under deep red light (720 nm) illumination to maintain dark adaptation. On the day of recordings, animals that had been dark-adapted overnight were anesthetized as described above and placed under a stereoscope on a regulated heating pad to regulate body temperature at 37°C. After inducing proptosis, methylcellulose was applied to the eye and an insulin needle was utilized to puncture the eye at the ora serrata. A 33-gauge metal tube was then inserted into the eye to deliver the solutions containing pharmacological agents. A 4 µl volume was injected into the eye at a rate of 50 nl s^-1^ by means of a foot-triggered pump (UltraMicroPumpIII, World Precision Instruments). The volume of the mouse vitreous has been reported to be 4.4 ± 0.7 µl^17^, so a near-total exchange of the vitreous was expected. A micropuncture was made at the top of the cornea to relieve intraocular pressure during the injection. After injection, animals were allowed to recover for 10 minutes in the dark before ERG recordings commenced.

### Pharmacology

Stock solutions of pharmacological agents were diluted into ultrapure, molecular biology grade phosphate-buffered saline (PBS, Affymetrix/USB™) containing 8 mM sodium phosphate, 1.9 mM potassium phosphate, 2.7 mM KCl and 137 mM NaCl. The solutions were titrated to pH 7.3 with aliquots of 10 M NaOH. Solutions were filtered with a 0.22 µm pore syringe PES filter (Olympus plastics, Genesee Scientific Corporation), split into 50 µl aliquots, and stored at 4°C, and used within a week of preparation.

#### AP4, PDA, Chloroquine, and Quinacrine

DL-AP4 sodium salt (Abcam Biochemicals), 2,3 cis-Piperidine dicarboxylic acid (Abcam Biochemicals), Chloroquine phosphate (Sigma-Aldrich), and Quinacrine dihydrochloride (Sigma-Aldrich) were all prepared following the same procedure. Each drug was diluted in PBS to obtain a 100 mM concentration. NaOH was then added to titrate the solution’s pH to 7.3. These drugs were use at 30 mM concentration.

#### VU590

VU590 hydrochloride (TOCRIS Bioscience) was dissolved in PBS, the pH adjusted to 7.3 with NaOH, and the solution left overnight at 4°C to allow the drug to fully solubilize. Once the solutions were clear with no precipitate, they were filtered and split into 50 µl aliquots. The VU590 concentration in intraocular injections was 1 mM.

#### Pentamidine

Pentamidine isethionate salt (Sigma Aldrich) was diluted in ddH_2_O to obtain a 100 mM concentration. This stock solution was then diluted into PBS to obtain the desired concentration. This drug was used at 1 mM concentration.

### Preliminary testing of dose-dependence

Preliminary experiments were performed to determine the concentrations in the 4 µl volume injected into the vitreous of AP4, PDA, chloroquine, pentamidine, quinacrine, and VU590 that elicited stable, maximal effects for the duration (up to 80 min) of an experiment. Information about the drugs’ targets and pertinent references is provided in **Table 1**.

**Table 1.** Pharmacological blocking drugs injected intravitreally and their target receptors.

| Drugs | Molecular target | Kir2.1; Kir4.1; Kir7.1<br>(IC<sub>50</sub>, μM) | 4 mL injection<br>(mM) | Figures* | Retina<br>refs. |
| --- | --- | --- | --- | --- | --- |
| L-AP4 (L-2-amino-4-phosphonobutanoic acid) | mGluR6 (ON-bipolar cells) | NKE | 30 | 3C, 3D; 6A-E | (a) |
| PDA (cis-2,3-piperidinedicarboxylic acid) | iGluRs (OFF-bipolar cells) | NKE | 30 | 3C, 3D; 6A-E | (b) |
| Chloroquine | Kir4.1 (Müller cells) | 8<sup>1</sup>, 1.2<sup>2</sup>; (0.5)<sup>3</sup>; NA | 30 | 4B, 4C | NA |
| Pentamidine | Kir2.1(Müller cells) | 0.17<sup>4</sup>; 0.097<sup>5</sup>; NA | 30 | 4B, 4C | NA |
| Quinacrine | Kir4.1 (Müller cells) | 65<sup>6</sup>; 1.8<sup>7</sup>; NA | 30 | 4B, 4C; 5A-D | NA |
| VU590 | Kir7.1 (RPE cells) | 10(NME); 10(NME); 8<sup>8</sup> | 1 | 4B, 4C | (c) |

*Abbreviations:* NKE, no known effect; NA, not available; (x)NME, no measureable effect at the concentration in parentheses.

## THEORY

The primary goals of the theoretical model developed herein are to predict the changes in the mouse subretinal space (SRS) of [K^+^] (hereafter, *K*_o, SRS_ or simply *K*_o_) that occur during rod light responses on the basis of the ionic mechanisms of rods (Fortenbach et al., 2021) (Peinado Allina, 2025b) and the RPE (Bialek and Miller, 1994), and to test the Oakley-Green-Steinberg-Miller (OGSM) hypothesis that these *K*_o_ changes give rise to a trans-epithelial potential of the retinal pigment epithelium (RPE) with the sign, kinetics and amplitude of measured *c*-waves.

### Model of the subretinal space (SRS)

#### Physical dimensions of the mouse SRS

Following (Steinberg, 1985a; Steinberg, 1985b), the SRS is defined as the contiguous volume of extracellular space (ECS) bounded on the posterior side by the apical membrane of the RPE, and on the anterior side by the external limiting membrane (ELM). The dimensions of the SRS were estimated from optical coherence tomography of live C57Bl6 mice and electron micrographs of fixed retina cut perpendicularly to the longitudinal axis of photoreceptor cells (Peinado Allina, 2025b), Figs. 1, 3). The axial length (*L*_SRS_) of the mouse SRS is ∼35 μm, but the apparent fraction of the retinal cross-sectional area occupied by the extracellular space *α*_*ECS*_ (*x*) varies with depth: based on our EM measurements it averages 9.5% ± 1.4% (mean ± s.d., n = 7) in the outer segment layer and 3.2% ± 1.4% (n = 12) in the inner segment layer. These estimates of *α*_*ECS*_ (*x*) are taken as upper limits, however, as abundant apical processes of Müller cells extend into the proximal SRS and of RPE cells into the distal SRS. Such processes not readily stabilized by fixation, but can be seen in optimally prepared cryosections –e.g., (Nawrot et al., 2004; Daniele et al., 2007a).

**Figure 1.**
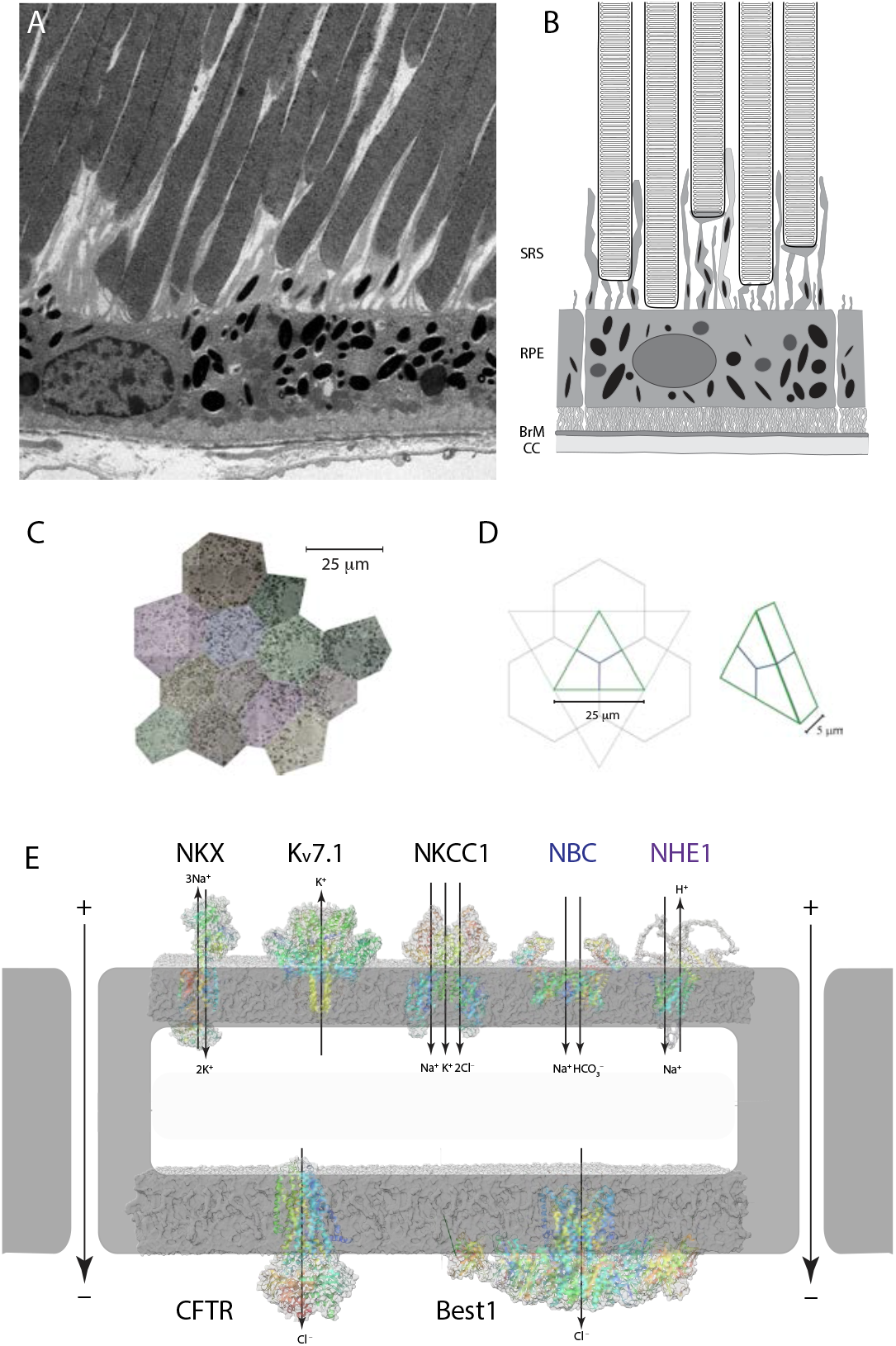
Anatomical and molecular characterization of the retinal pigment epithelium (RPE). **A**. Electron micrograph of the posterior retina of a C57Bl/6 mouse with rod outer segments and an RPE cell. **B**. Schematic drawing comparable to panel A, with components labeled (CC, choriocapillaris; BrM, Bruch’s membrane; SRS, sub-retinal space). The SRS extends from the apical RPE to the external limiting membrane (not shown); numerous Müller cell microvilli expressing Kir7.1 project into the SRS. **C**. *En face* electron micrograph composite of a portion of mouse RPE, colorized to segment the 12 cells. **D**. Unit cell of a model RPE with hexagonal packing: given 5 μm thickness and an average geometrical surface area of 300 μm^2^ the intercellular gap width is calculated to be 3 nm. **E**. Molecular components of the apical and basal RPE membrane that maintain the cell’s polarity, giving rise to a paracellular current that flows from the apical to basal side (larger arrows in the gaps). The contributions of these components to the resting transepithelial potential and its change when SRS *K*_o_ declines during the light response are quantified in the model of the mouse RPE (Eqs 10 – 23; Supplementary Information, “3. Kinetic model of the RPE response to changes in SRS *K*_o_”)

#### Diffusion of K^+^ in the SRS

Unimpeded diffusion of K^+^ in tissue has been estimated to have a coefficient of *D*_K_ = 2400 μm^2^ s^-1^ (cf 5 studies cited in (Park and Durand, 2006). Thus, if axial diffusion of K^+^ in the SRS is unimpeded, equilibration between the ELM and RPE apical surface should occur on the scale of *t*_equil_ = *L*^2^ _SRS_ / (4*D*_K_ ) < 0.2 s. Müller cell and RPE apical processes extending into the SRS affect not only its volume, but also slow diffusion by increasing tortuosity (*λ*), estimated to reduce *D*_K_ by a factor *λ*^2^ ∼ 4 in amphibian retina (Newman and Odette, 1984). Consequently, diffusional mixing of K^+^ in the SRS may need to be considered for theory applied on times scales of the order of 1 s. We implemented SRS K^+^ diffusion as an option in the model, using the formalism of (Calvert et al., 2010), which treats 1D (axial) diffusion in contiguous compartments of differing cross-sections. However, the “well-stirred” variant of the SRS model was used for the results presented herein.

#### K^+^ fluxes and the dark (resting) steady state of SRS K_o_ in a “well-stirred” SRS

A necessary condition for rods, RPE and the SRS to be in steady state is that the concentration *K*_o, SRS_ be constant. Fluxes of K^+^ to/from the SRS comprise transmembrane fluxes from rod inner and outer segments (*φ*_rods_ ), RPE cells’ apical membranes (*φ*_RPE_ ) and Müller cell processes extending into the SRS (*φ*_MC_ ), and fluxes across the external limiting membrane (*φ*_ELM_ ) and the RPE paracellular pathway (*φ*_RPE, PC_ ). These fluxes were expressed in common units (mol cm^-2^ s^-1^), and in the resting (dark adapted) steady state must sum to zero:

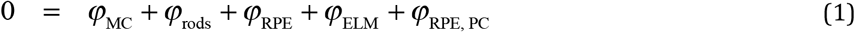

The transmembrane fluxes are further elaborated below in terms of the contributions from the specific underlying ionic mechanisms: rods, NKX (α3β2 isoform), Kv2.1 and HCN1 channels; RPE cells, Kir7.1 and NKX (α1β1 isoform); Müller cells (MC), Kir4.1 channels. The units follow from treating the retina as a series of transverse slabs of defined thickness: the rod current or K^+^ flux density at depth *x* is that of a single rod at that depth multiplied by the rod density, and similarly for the Müller cells. RPE currents and apical K^+^ fluxes are naturally expressed as densities with respect to the slab or node of the spatial origin (*x* =0)

The mouse ELM is about 1 μm thick (Peinado Allina, 2025b, Fig. 1), and is formed by densely packed Müller cell apical processes interconnected with each other and photoreceptor cells by adherens junctions (Daniele et al., 2007). We assume that the ELM acts as a barrier to ion movement, but that ECS fluxes across it obeys Fick’s Law, so that the trans-ELM K^+^ flux is given by

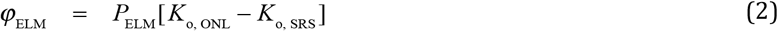

where *P*_ELM_ (cm s^-1^) is the permeability coefficient of the ELM for K^+^, and *K*_o, ONL_ is the concentration in the outer nuclear layer, and for unit consistency the concentrations *K*_o, ONL_ and *K*_o, SRS_ are expressed in mol cm^-3^. Calculation of K^+^ flux through the RPE paracellular pathway must take into consideration the resting transepithelial potential, which renders the SRS at the RPE apical face positive by 5 to 10 mV relative to the basolateral ECS. The trans-epithelial barrier can be treated with the Nernst-Planck formalism (la Cour et al., 1986), yielding the relation

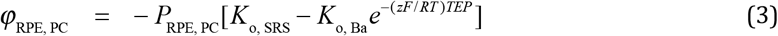

where *P*_RPE, PC_ (cm s^-1^) is the permeability coefficient of the RPE paracellular path for K^+^, and *K*_o, Ba_ the K^+^ concentration in the ECS on the basal side of the RPE; this term represents efflux of K^+^ from the SRS when *K*_o, SRS_ > *K*_o, Ba_ *e*^−( *zF* / *RT* )*TEP*^ . By dint of the negligible reflection coefficient (high permeability) for K^+^ of the vascular endothelial cell membrane (Renken and Curry, 1978; Michel, 2013), the close proximity of the rich choroidal vasculature to the basal RPE, and of the extensive vasculature in the outer plexiform layer immediately proximal to the ELM, we assume that *K*_o, ONL_ = *K*_o, Ba_ = *K*_o, plasma_ (**Fig. 2B**) Equations (1) – (3) can be solved for *K*_o, SRS_, yielding

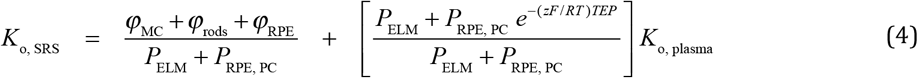

**Figure 2.**
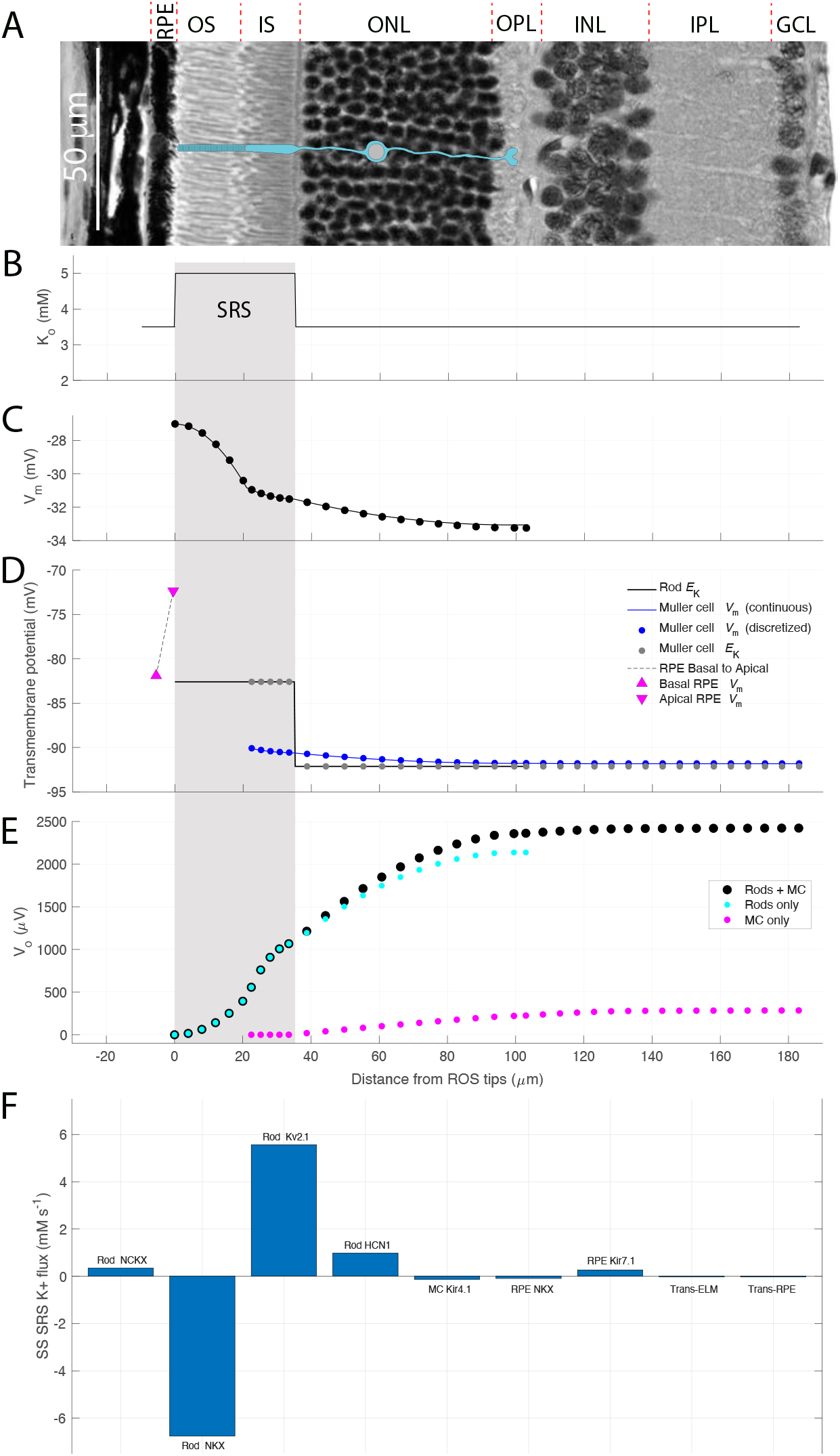
K^+^ fluxes to the subretinal space in the dark steady-state are dominated by fluxes of NKX transporters and K_v_2.1 channels expressed in rod inner segments. **A**. Image of plastic section of C57Bl/6 central retina, with layers identified above (RPE, retinal pigment epithelium; OS, outer segment layer; IS, inner segment layer; ONL, outer nuclear layer; OPL, outer plexiform layer; INL, inner nuclear layer; IPL, inner plexiform layer; GCL, ganglion cell layer). A schematic rod (light blue) has been overlaid on the section. **B**. Distribution of [K^+^]_o_ in the dark, resting state. The light gray bar highlights the subretinal space (SRS). **C**. Resting membrane potential distribution of the average rod of the 11-rod ensemble model (cf (Peinado Allina, 2025a) Fig. 5A). **D**. Distributions of K^+^ equilibrium potentials of RPE cells (symbols), rods (black line) and Müller cells (gray dots), and the Müller cell membrane potential (blue line; continuous solution; dots, discretized solution). (Rods and Müller cells are assumed to have the same [K^+^]_I_, 110 mM. The electrochemical driving force for K^+^ for Müller cells is inward in the SRS and outward elsewhere in the retina, leading to a circulating current that sinks in the portion of the Müller cell in the SRS and has its sources distributed throughout the rest of the cell (cf. Appendix II). **E**. Extracellular potentials arising from the rod (cyan dots) and Müller cell (magenta dots) circulating currents. The black filled symbols plot the potential arising from the combined extracellular current of rods and Müller cells. **F**. Fluxes of K^+^ to/from the SRS from cellular mechanisms and trans-layer transport (cf. Eqs 1 – 3). The fluxes are expressed in SRS molar units, and sum (Eq 1) approximately to zero (< 0.008 mM s^-1^). The SRS volume density (*V*_SRS_ in Eq 5) used in calculating the molar fluxes was 0.755 μL cm^-2^.

**Figure 3.**
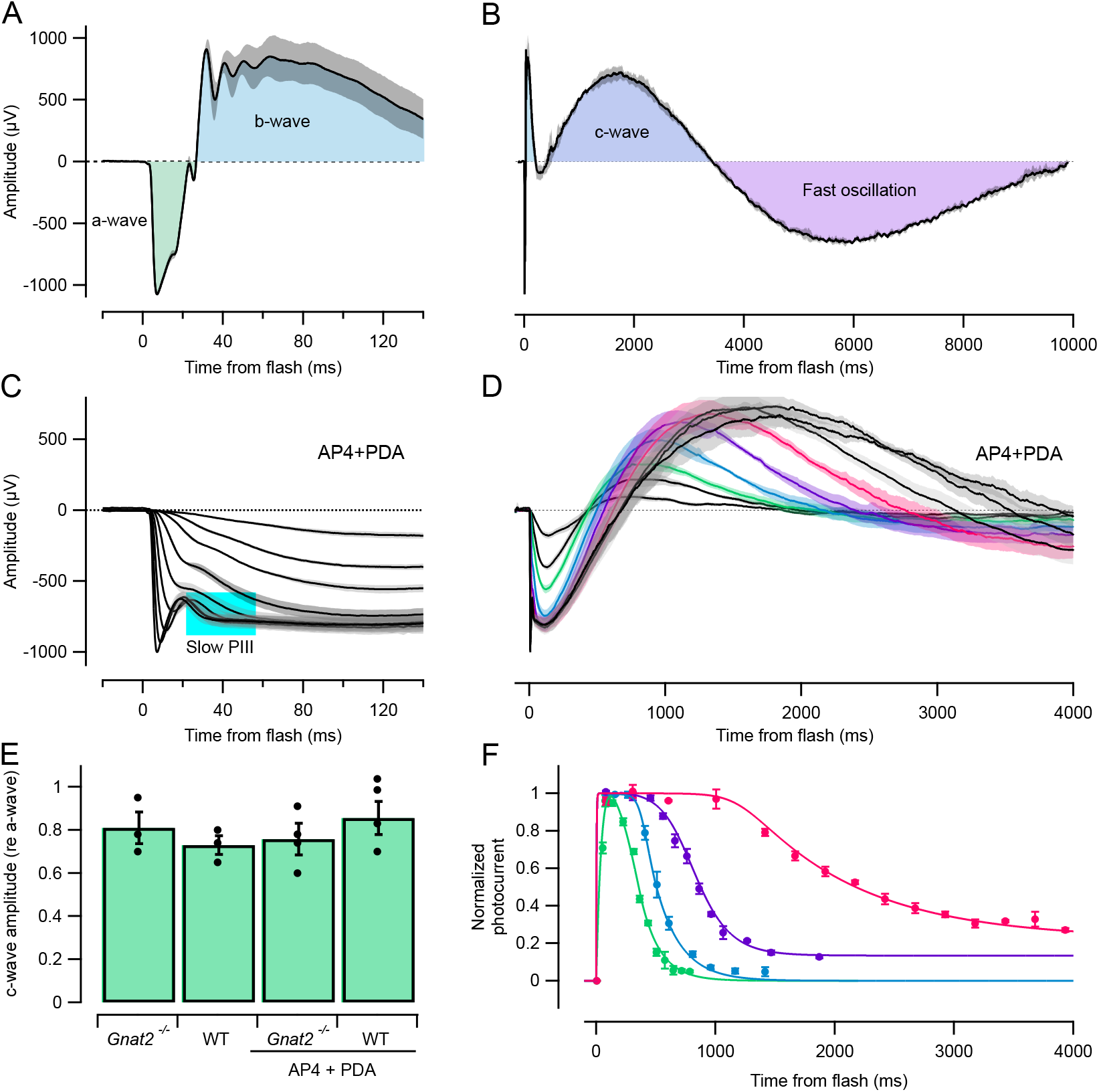
Characteristics of rod-driven *a*- and *c*-waves after intraocular injection of blockers of glutamatergic metabotropic (AP4) and ionotropic (PDA) synaptic transmission. **A, B**. Averaged ERG of a C57Bl/6 mouse to a 500 nm flash delivering 5×10^5^ photons μm^-2^, with identification of the principal standard components; the same data traces are plotted on two different abscissas in the two panels to reveal the time scales of the components. **C, D**. Family of ERGs obtained 30 min after intraocular injection of the glutamatergic synaptic transmission blockers AP4 and PDA. The early time window (C), reveals the positive-going *b*-wave to be completely abolished, while the longer time window prominently shows the corneal-positive *c*-waves. **E**. Bar chart of saturated *c*-wave amplitudes for the two genotypes (*Gnat2*^−/−^ and WT (C57Bl/6) controls; *c*-waves were measured from the baseline and scaled relative to the amplitude of the saturated *a*-wave obtained in the same experiment. (The unscaled amplitudes are presented in Table 3.) **F**. Recovery of rod photoresponses of WT mice measured with a paired-flash protocol (replotted from (Peinado Allina et al., 2017)). The colorization in F corresponds to that in panel D, and serves to illustrate the status of the rod circulating current during the *c*-waves (note the identical time bases). The *c*-wave model (**Fig. 7**) was applied to the average responses in D to the two most intense stimuli (black traces).

Examination of the terms on the right-hand side of Eq 4 helps explain several features of the regulation of *K*_o, SRS._ Firstly, the second term on the right-hand side can approach but must be lower than *K*_o, plasma_. Secondly, the first term on the right-hand side shows that steady-state *K*_o, SRS_ can exceed or be below *K*_o, plasma,_ depending on whether cytosolic K^+^ uptake by the NKX’s of the rods and RPE exceeds K^+^ efflux through other membrane ionic mechanisms. As analysis of the transcellular K^+^ fluxes (RESULTS) indicates the numerator of this term to be positive at rest, achievement of a dark steady-state necessitates K^+^ efflux across the two extracellular barriers (Eqs 2 –3), and so explains why resting *K*_o, SRS_ exceeds *K*_o, plasma_. Third, Eqs 1 -4 yield a rate equation governing *K*_o, SRS_:

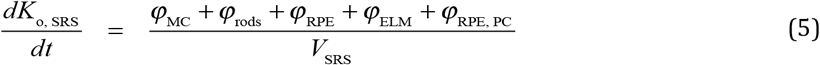

As the terms in the numerator of Eq 5 have been expressed in flux density units (mol cm^-2^ s^-1^), the denominator *V*_SRS_ must be expressed as a “volume density” (units: L cm^-2^), so that both sides of the equation will have the units of M s^-1^. For*α*_*ECS*_ (*x*) as described above, an initial estimate from analysis of EM data gave *V*_SRS_ = 0.23 μL cm^-2^. Fourth, given specific values for *K*_o, SRS_ and *K*_o, plasma_ and the cellular K^+^ flux terms, Eq 4 can be solved for *P*_ELM_/*P*_RPE,PC_, providing a useful constraint in the estimation and assessment of parameters.

In the following sections we provide physiological details of the cellular trans-membrane flux terms and estimates of their values, noting here that several of the terms are functions of *K*_o, SRS_. Thus, K^+^ fluxes through rod K_v_2.1 channels, and through RPE Kir7.1 and Müller cell Kir4.1 channels depend on *E*_K_ at their respective membranes facing the SRS, which in turn depends on *K*_o, SRS_ through the cells’ Nernst potentials. In addition, both rod and RPE NKX activities are weakly dependent on *K*_o, SRS_ through binding interactions (Stanley et al., 2015), as detailed below.

### Model of rod photoreceptor K^+^ flux to/from the SRS

#### Electrical model of the rod

The outer and inner segments were assumed to be identical for all rods, as in the companion paper; including the identities, distributions and other properties of the CNG channels and NCKX of the outer segment, the K_v_2.1 channels and the NKX of the inner segment membrane and the HCN1 channels (Peinado Allina, 2025b). For key conclusions based on the theory, the rod membrane potential was in one of two steady states: the dark adapted state, or the light-exposed state in which the CNG-gated channels are completely closed. While not isopotential in the dark adapted state due to the flow of the dark current through the axial resistance of the outer segment, the rod membrane potential is essentially constant over the inner segment (Peinado Allina, 2025b) and taken to be −32 mV in darkness, as measured in mouse retinal slices (Fortenbach et al., 2021). The membrane potential predicted for the light-saturated steady state (after the HCN1 “nose” relaxation) was −48 mV to −47 mV (**Fig. S1A**).

#### Fluxes of K^+^ and Na^+^ across the rod plasma membrane in the dark

In the dark steady state, rods must maintain homeostasis of all permeant ions. The fluxes of K^+^ and Na^+^ through the five outer and inner segment ionic mechanisms were calculated as described in (Fortenbach et al., 2021) and (Peinado Allina, 2025b), but is recapped here, as follows. In the dark adapted state Na^+^ enters the rod through the CNG channels and the NCKX in the outer segment, and so the quantity of Na^+^ influx is determined, given a total inward dark current of 20 pA. As the NKX of the inner segment is the only mechanism of Na^+^ efflux from rods, its electrogenic current in darkness is readily determined to be 6.8 pA (34%), and given the 3:2 Na^+^:K^+^ stoichiometry of the NKX, the dark K^+^ influx is 1.4 fmol s^-1^ rod^-1^. In the dark this K^+^ influx into the rod is balanced by an equal efflux, primarily through K_v_2.1 channels (Fortenbach et al., 2021) (**Fig. 2F**).

#### Fluxes of K^+^ and Na^+^ across the rod membrane in the light-stimulated state

With all CNG channels closed by a strong light exposure, the outer segment has negligible ionic flux on the time scale of the *c*-wave, as the NCKX transporter drives Ca^2+^_i_ to a minimum in less than 150 ms (Fortenbach et al., 2021). However, even after the light-exposed rod hyperpolarizes to an essentially constant level, ionic fluxes across the inner segment membrane can continue. A primary mechanism of such ion flux is the NKX, which will continue to transport K^+^ into the rod and export Na^+^ to the SRS at a rate similar to that exhibited in the dark state. The electrogenic current of the NKX follows the relation

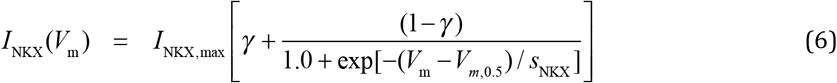

The dependence of the NKX current on *K*_o_ in Eq 6 is from (Stanley et al., 2015), while the Boltzmann function is that previously used to describe the currents of the α3β2 (rod) isoform by (Fortenbach et al., 2021). Based on the 3Na^+^ out:2 K^+^ in stoichiometry of the NKX, the SRS flux of K^+^ owing to the rod NKX is given by

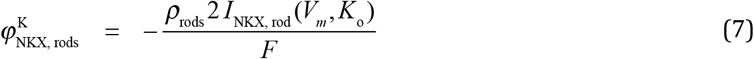

where *F* is the Faraday constant.

A second mechanism of rod K^+^ and Na^+^ transmembrane flux in the light-exposed state is the HCN1 channels, whose reversal potential (*E*_H_ = −31 mV) reveals them to be permeant to both cations. Though rod HCN1 channels carry negligible current in the dark (Fortenbach et al., 2021), in the light-exposed, hyperpolarized state their inward current is predicted to be −9 pA, depolarizing the rod from its peak hyperpolarization, and contributing to the balancing of the outward electrogenic NKX current (Peinado Allina, 2025b, Fig. 7E) . The HCN1 current is describable with 2-state Boltzmann kinetics whose steady-state current for the membrane potential *V*_m_ (Fortenbach et al., 2021) obeys the relation

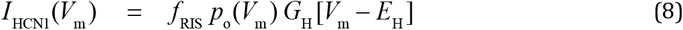

where *f*_RIS_ = 0.75 is the fraction of HCN1 expressed in the rod inner segment, *p*_*o*_(*V*_m_) is the open probability, *G*_H_= 1.4 nS is the total conductance, and *E*_H_ the reversal potential. To quantify K^+^ and Na^+^ fluxes through HCN1 channels we equated Eq 8 with the Goldman-Hodgkin-Katz current relation for a conductance permeable to both Na^+^ and K^+^ (Lewis, 1979) at *V*_m_ = *V*_m, Ap, rest_ :

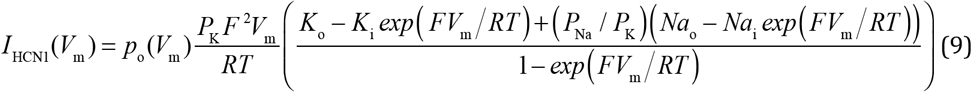

Dividing Eq 9 by the Faraday constant and extracting the term for K^+^ gives the corresponding K^+^ flux (mol s^-1^ rod^-1^). (A parallel relation holds for the flux of Na^+^.) While Eq 9 provides a means of estimating the fluxes of the two HCN1-permeant cations, it is problematic inasmuch as the predicted current does not generally agree with the measured voltage-dependence (Eq 8). To obviate this problem we equated Eq 9 with Eq 8 at each *V*_m_ and thereby extracted component K^+^ and Na^+^ fluxes (cf. Supplementary Information, “1. K^+^ flux through mouse HCN1 channels”).

#### Mouse rod responses to flashes that produce maximal c-waves

The recovery of the mouse rod photoresponse for flash strengths exceeding ∼ 1 photoisomerization per disc face (∼ 1500 isomerizations for a 20 μm outer segment) slows down substantially from that which governs the responses at lower intensities (Lyubarsky and Pugh, 1996; Martemyanov et al., 2008), and a full family of flash-activated *c*-waves is clearly observed only in response to such intensities. To characterize the recovery of mouse rods at the high intensities that produce maximal *c*-waves, we used mouse rod photocurrent responses extracted from ERG *a*-waves with the paired-flash method (Peinado Allina et al., 2017, Fig. 3F) (cf “Supplementary Information, “2. Mouse rod responses *in vivo* to stimuli that generate saturating *c*-waves”).

### Electrophysiological model of the retinal pigment epithelium (RPE)

#### Overview

The Oakley-Green-Steinberg-Miller (OGSM) hypothesis linking light-stimulated changes in SRS *K*_o_ to a change in RPE transepithelial potential (TEP) postulates that decrease in *K*_o_ caused by continued inward pumping by rod NKX in the presence of reduced outward K_v_2.1 current causes a negative shift in the K^+^ Nernst potential of the RPE apical membrane. The shift in the K^+^ Nernst potential in turn causes an increase in outward current through the abundant Kir7.1 channels in the RPE apical membrane, increasing the outward current that must sink across the basolateral membrane through Cl^−^ channels **(Fig. 1E**), passing from apical to basal membrane through the paracellular path. The flow through the resistive paracellular path causes a retinal-positive” TEP, manifest as a corneal-positive potential identified as the *c*-wave (Oakley and Green, 1976; Oakley, 1977; Steinberg et al., 1980). The model of the RPE developed here is grounded in the foundational work of Oakley, Green Steinberg, Miller and colleagues, but includes quantitative information from the mouse about the kinetics and magnitude of the light-stimulated decrease in SRS *K*_o_, and specification of the molecular identities (**Fig. 1E**) and properties of key ionic mechanisms not available when much of the seminal research was done.

#### Structure and transepithelial resistance

The murine RPE comprises an ∼ 5 μm thick monolayer of approximately hexagonally packed cells interlinked by tight junctions. The cells are approximately uniform in size across the retina, and have an average cross-sectional area of ∼300 μm^2^ (Kim et al., 2021). The average resistance of mouse RPE cell layers in primary culture is 200–400 Ω cm^2^ (Fernandez-Godino et al., 2016).

#### Molecular identities of the principal RPE ion channels and transporters

Based on the extensive physiological investigations of Miller and colleagues of mammalian RPE (Bialek and Miller, 1994; Gallemore et al., 1997; Adijanto et al., 2009; Maminishkis and Miller, 2010; Miyagishima et al., 2017) we assumed the primary ionic mechanisms determining the resting state of the RPE cell to be those shown schematically in **Fig. 1F**.

#### Ionic mechanisms of the RPE apical membrane

Electrical activity of the RPE apical membrane is dominated by two ionic mechanisms: NKX (primarily α1β1 with some α1β2) and Kir7.1 (Kcnj13); (Shimura et al., 2001; Cornejo et al., 2018). The dependence of the electrogenic current of various NKX isoforms on membrane potential and *K*_o_ has been described in detail by (Stanley et al., 2015), from whose work we developed the following representation of the NKX current density (cf (Fortenbach et al., 2021):

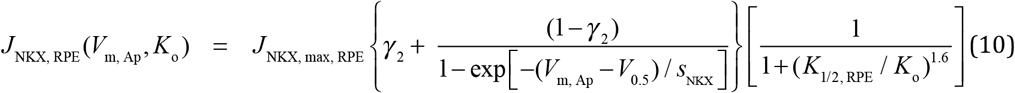

Here *J*_NKX, max, RPE_ (A cm^-2^) is the maximum NKX current, γ_2_ = 0.38, *V*_0.5_ = -71 mV, *s*_NKX_ = 23.2 mV, and *K*_1/2, RPE_ = 1.8 mM.

The electrical properties of RPE Kir7.1 currents have been extensively characterized by (Shimura et al., 2001), who reported that they accounted for most of the K^+^ conductance of isolated bovine RPE cells, and also showed, as have others (Yang et al., 2003), that Kir7.1 expression in the RPE is confined to the apical membrane. Based on Shimura et al’s whole-cell currents we characterized the Kir7.1 current density of the apical RPE with the following Boltzmann relation:

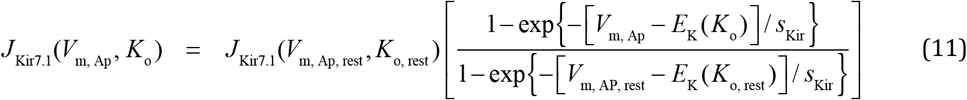

Here, *J*_Kir_ (*V*_m, Ap, rest_, *K*_o, rest_ ) (A cm^-2)^ is the outward current at the resting apical membrane potential, and the numerator term in the rectangular bracket gives the Boltzmann dependence on apical membrane potential V_m, Ap_ and on *E*_K_(*K*_o_), the Nernst potential for an SRS potassium concentration *K*_o_, and *s*_Kir_ is a steepness factor. The term in the denominator of the rectangular bracket of Eq 9 is a fixed scaling factor. (A graphical presentation of Eq 11, along with the fitting of the Boltzmann relation to the data of (Shimura et al., 2001) is provided in the Supplementary Material, “3. Kinetic model of the RPE response to changes in SRS *K*_o_”).

#### Ionic mechanisms of the RPE basolateral membrane

Electrogenic NKX activity and Kir7.1 outward current is balanced at rest by inward current in the basolateral membrane (Bialek and Miller, 1994). This spatial separation of the outward and inward currents causes current to flow extracellularly through the paracellular pathway *en route* to the basolateral sink. (This extracellular circulating current is similar to that created by the photoreceptors’ dark current, and indeed the paracellular component of the RPE circulating current flows in the same direction as the photoreceptor dark current.) The identity of all ionic mechanisms carrying the resting inward current in mouse have not been definitively established, but the basolateral membrane very highly expresses bestrophin (BEST1), a tetrameric chloride channel (Sun et al., 2002; Bakall et al., 2003; Qu et al., 2004; Owji et al., 2020; Owji et al., 2021), and is also known to express CFTR (Liu et al., 2012). The current/voltage (I/V) relation of mouse Best1 is essentially linear (Marmorstein et al., 2015), and so we described the basolateral chloride current as follows:

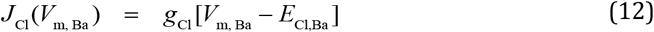

The chloride concentration *Cl*_i_ in resting bovine RPE cells in medium with *K*_o_ = 5 mM has been measured to be 65 ± 10 mM (Bialek and Miller, 1994). Given the very close proximity of the RPE basal membrane to the choriocapillaris, and that C57Bl6 serum chloride is 107-111 mM (Otto et al., 2016), *E*_Cl, Ba_ = −14 mV. As resting V_m, Ba_ is ∼ −70 mV (Bialek and Miller, 1994), Eq 12 predicts a resting inward current, which we take to be equal in magnitude to the resting outward current of the apical and basal membranes, which determines *g*_Cl_.

The basal RPE is also known to have K^+^ conductance (Bialek and Miller, 1994) whose molecular identity has not yet been determined. We assumed this current to obey a linear I/V relation:

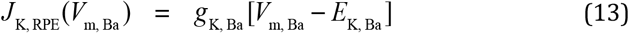

Equations 10 −13 define the properties the electrical mechanisms of the RPE model. The RPE membranes also contains a number of non-electrogenic transport mechanisms (Bialek and Miller, 1994; Adijanto et al., 2009): three of these (NKCC, NBC, NHE) are essential for maintaining the resting flux balance of K^+^, Na^+^ and Cl^−^, and need to be included in defining the RPE resting state, as follows.

#### The resting state of the RPE specifies a unique set of membrane currents

Dark adaptation constitutes a special steady state for the retina, as night provides an extended stable ambient condition for photoreceptors and consequently other cells to come into both electrical balance and homeostatic equilibrium of permeant ions. The ionic currents and ion fluxes of an RPE cell comprising the membrane transporters NKX, NKCC, and channels permeable to K^+^ and Cl^−^ (Fig. 3E) are uniquely determined up to one scale constant. This result follows from the following conservation relations,

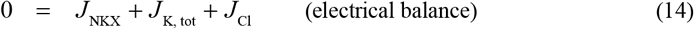

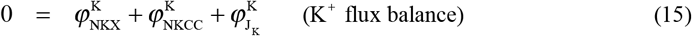

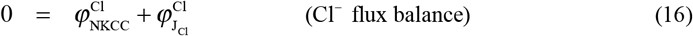

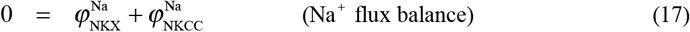

Six additional flux balance relations are dictated by the NKX and NKCC transporters’ stoichiometries, and the conversion of electrical currents to ion fluxes:

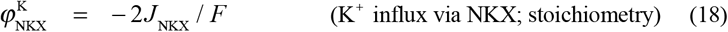

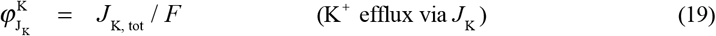

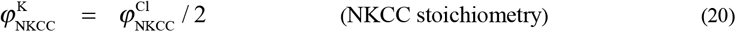

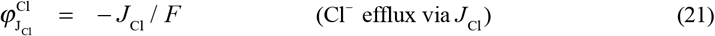

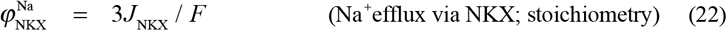

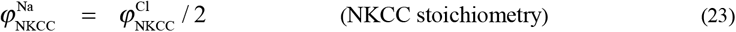

In Eqs 14-23 the symbol 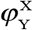 represents the transmembrane flux density (mol cm^-2^ s^-1^) of ions of type X via a channel or transporter of type Y. (Fluxes obey the same sign convention as positive ions: inward flux is negative, outward positive. However, the outward flux of negative Cl^−^ ions via chloride channels constitutes an inward electrical current.) In Eqs 14, and 19, *J*_K,tot_ represents the total K^+^ current density of both apical (Kir7.1, Eq 11) and basolateral (Eq 13) membranes. Equations 14−23 comprise a system of 10 homogeneous linear equations in 10 unknowns, whose solution is unique up to a scaling constant that can be conveniently taken to be the current density of the RPE NKX, *J*_NKX_ (A cm^-2^): thus, the other nine variables can be expressed as linear multiples of resting *J*_NKX_ and derived analytically (Appendix). The equations embody the constraint *J*_K, tot_ = 5 *J*_NKX_ . Resting *J*_NKX_ was initially assigned the value 2.5 μA cm^-2^ taken (Shimura et al., 1999), but was treated as a potentially variable parameter.

#### Resting trans-epithelial potential (TEP)

The resting currents *J*_NKX_ and *J*_Kir7.1_ are outward (source) currents that must be balanced by inward current through basolateral chloride channels (cf **Fig. 3E**). The standard convention for the ERG implicitly defines extracellular current flowing from the anterior to the posterior of the retina as negative (i.e., the *x-*axis increases from the posterior to the anterior of the eye): thus, for example, the extracellular rod dark current is negative, flowing “downhill” its ECS potential toward outer segment sinks (Fig. 4E). For consistency with this convention, the TEP of the RPE is defined as the potential difference between the apical (anterior) and basal (posterior) extracellular space, which in turn can be related to the apical and basal membrane potentials as follows:

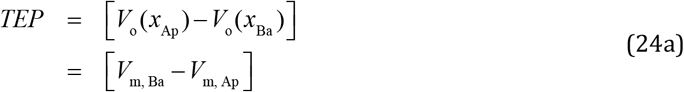

**Figure 4.**
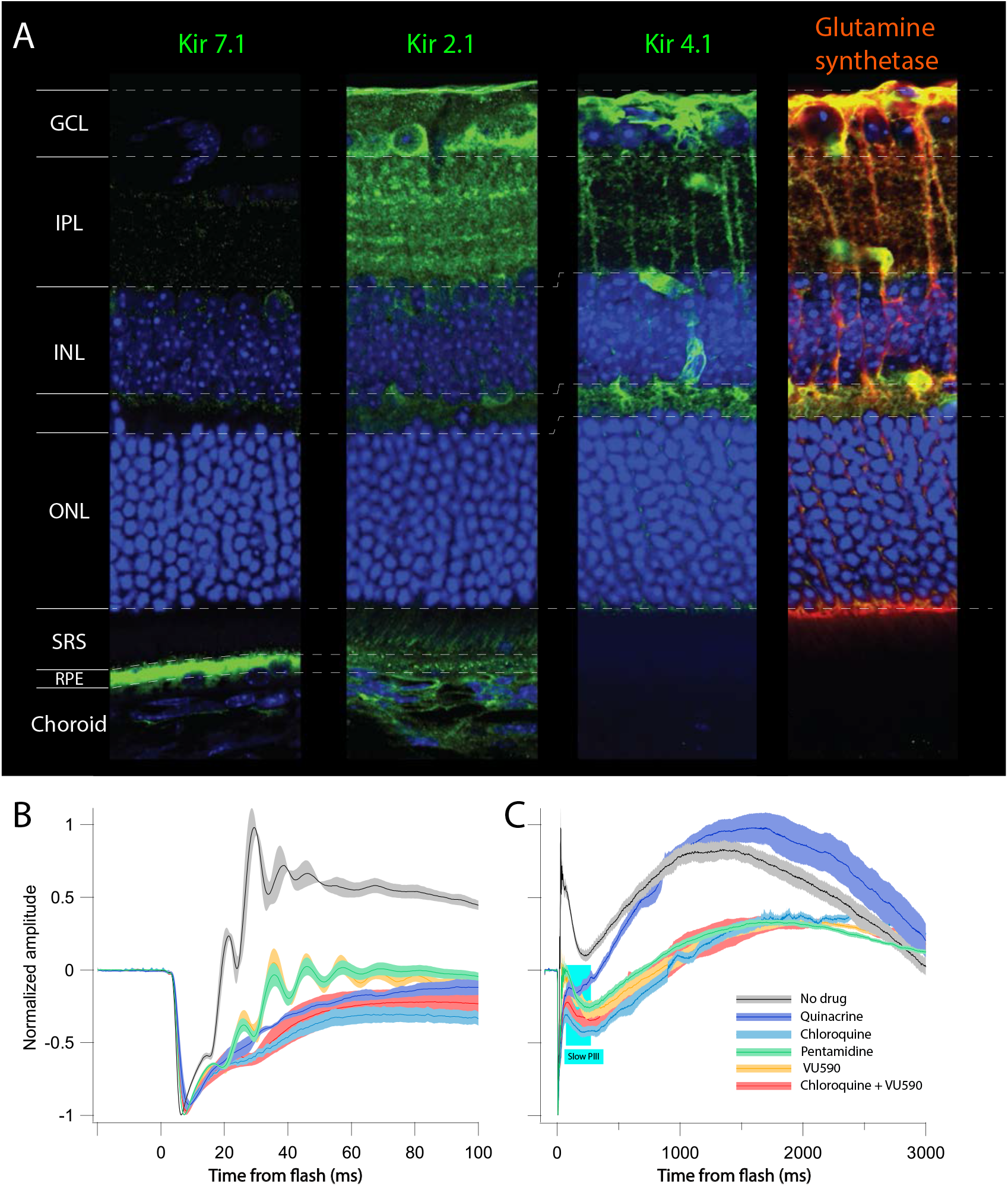
Immunohistochemical localization in the mouse retina of Kir channels and ERGs measured after intravitreal injections of Kir-blocking drugs. **A**. Immunofluorescence (IF) of cryosections stained with antibodies of three Kir channels expressed in the retina, and combined IF of glutamine synthetase (a Müller cell marker) and Kir4.1 **B, C**. Averaged ERGs obtained in response to a flash delivering 5×10^5^ photons μm^-2^ after intravitreal injections of Kir pore-blocking agents *(cf* Table 1) shown on two time scales, 100 ms (A) and 3 s (B). The traces were scaled to have unity magnitude *a*-waves (absolute amplitudes are provided in **Table 3**). In panel C a gray rectangle identifies the time-magnitude region of the negative-going “Slow PIII” component, which was observed in the presence of all drugs except quinacrine.

(The second line of Eq 24a can be derived by moving a unit charge *q* out the RPE apical membrane, through the paracellular path and back into the RPE cell interior through the basal membrane: as the work done must sum to zero, 0 = −*qV*_m, AP_ + *q* [*V*_o_ (*x*_Ba_ ) − *V*_o_ (*x*_Ap_ )] + *qV*_m, Ba_, which yields the second line of 23a.) The TEP must also satisfy Ohm’s current law:

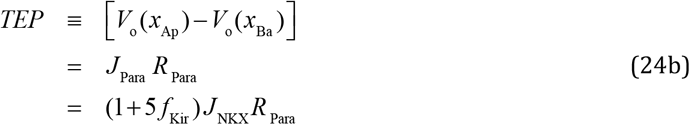

Where *J*_Para_ = −( *J*_NKX_ + *J*_Kir_ ), (A cm^2^) is the current dnsity, and *f*_Kir_ = *J*_Kir_ / *J*_K, tot_, is the fraction of total K^+^ current carried by the Kir7.1 channels, *R*_Para_ (Ω cm^2^) the paracellular resistance.

*Calculating the trans-RPE potential response to a transient change in SRS K*_o._ To predict the TEP arising from changes in SRS *K*_o_, the following pair of coupled differential equations for the apical and basal membrane potentials must be solved:

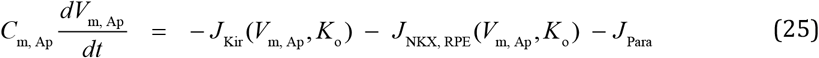

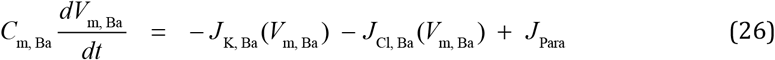

The forcing function for Eqs 25, 26 is the change in SRS *K*_o_ caused by rod light responses (Eqs 6–9). Insight into RPE dynamics and the role of Kir7.1 can be garnered by solving Eqs 25-26 for step or ramp changes in SRS *K*_o_ (SI, “3. Kinetic model of the RPE response to changes in SRS *K*_o_”).)

### Rest state and initial conditions of the dynamic model: constraints and parameters

Application of the model comprised two phases: (1) establishment of the dark steady-state for a specific set of parameters (**Table 2**); (2) solution of the set of ordinary differential equations (ODEs) governing the model variables during a rod light response (CNG channel closure). The dark steady-state analysis determined all initial conditions for the ODEs, and running the model in the absence of a light stimulus served as check of the initial conditions, requiring negligible drift in all variables.

**Table 2.** Variables and Parameters of the SRS –RPE Model.

| *Variable* | *Unit* | *Equation* | *Description/Comment* |
| --- | --- | --- | --- |
| $\phi_{\text{rods}}^{\text{K}}$ | $\text{mol cm}^{-2} \text{ s}^{-1}$ | 1 | Net SRS $\text{K}^+$ flux to/from rods |
| $\phi_{\text{RPE, Ap}}^{\text{K}}$ | $\text{mol cm}^{-2} \text{ s}^{-1}$ | 1 | Net SRS $\text{K}^+$ flux to/from apical RPE |
| $V_{\text{m, Ap}}$ | mV | 10, 11 | RPE apical membrane potential |
| $V_{\text{m, Ba}}$ | mV | 12, 13 | RPE basolateral membrane potential |
| $J_{\text{NKX, RPE}}$ | $\mu\text{A cm}^{-2}$ | 10 | RPE apical NKX electrogenic current |
| $J_{\text{Kir7.1}}$ | $\text{A cm}^{-2}$ | 11 | RPE apical Kir7.1 current |
| $J_{\text{Para}}$ | $\text{A cm}^{-2}$ | 24b, 25, 26 | RPE paracellular current |
| $J_{\text{Cl}}$ | $\text{A cm}^{-2}$ | 12 | RPE basolateral $\text{Cl}^-$ current |
| $J_{\text{K, Ba}}$ | $\text{A cm}^{-2}$ | 10 | RPE basolateral $\text{K}^+$ current |
| $K_{\text{o, SRS}}$ | mM | 1, 7, 8 | SRS concentration of $\text{K}^+$ |

| *Parameter* | *Unit* | *Equation* | *Range* | *Value* | *Description/Comment* |
| --- | --- | --- | --- | --- | --- |
| $R_{\text{Para}}$ | $\Omega \text{ cm}^2$ | 24b | 200 – 400 | 300 | RPE layer paracellular resistance |
| $V_{\text{m, Ap, Rest}}$ | mV | | NA | – 69 (fixed) | Resting RPE apical membrane potential |
| $V_{\text{m, Ba, Rest}}$ | mV | | NA | – 62.5 (fixed) | Resting RPE basal membrane potential |
| $J_{\text{NKX, RPE, rest}}$ | $\mu\text{A cm}^{-2}$ | | 2.5 – 7.0 | 4.0 | RPE apical membrane NKX current density |
| $g_{\text{Cl, Ba}}$ | $\text{S cm}^{-2}$ | | (EOC) | $6.2 \times 10^{-4}$ | RPE basal chloride conductance |
| $K_{\text{o, Plasma}}$ | mM | 4 | NA | 3.5 (fixed) | $[\text{K}^+]$ in mouse blood plasma |
| $K_{\text{o, SRS, rest}}$ | mM | 4 | NA | 5.0 (fixed) | $[\text{K}^+]$ in the SRS in dark adapted steady-state |
| $P_{\text{ELM}}$ | $\text{cm s}^{-1}$ | 2 | $10^{-5} - 10^{-4}$ | $2.0 \times 10^{-5}$ | $\text{K}^+$ permeability of the ELM |
| $P_{\text{RPE}}$ | $\text{cm s}^{-1}$ | 3 | $10^{-5} - 10^{-4}$ | $1.35 \times 10^{-5}$ | $\text{K}^+$ permeability of the RPE layer |
| $V_{\text{SRS}}$ | $\text{L cm}^{-2}$ | 5 | $3 \times 10^{-7} - 2 \times 10^{-6}$ | $7.55 \times 10^{-7}$ | Volume density of the SRS |

indicates not applicable because the parameter was fixed, while EOC” indicates the parameter was set in satisfying the Equation of Continuity.) The table focuses on variables and parameters unique to the paper; those describing the rod ensemble elements of the model are given in (Peinado Allina, 2025b), though occasionally repeated herein for expositional clarity of the ionic mechanisms contributing to SRS $K^+$ flux. A dynamic model of the Müller cell was not achieved, but a steady-state model was developed and an empirical description of the light-driven source in the absence of glutamatergic synaptic transmission was obtained with quinacrine pharmacology (cf. Supplementary Information, “4. Electrophysiological Model of the Mouse Müller Cell and the Extraction of Slow PIII”).

**Table 3.** Effects of pharmacological agents on the magnitudes of mouse ERG *a*- and *c*-waves.

| Genotype | Drug | *a*-wave (μV) | *c*-wave (μV) | *c*-/*a*-wave | Figures |
| --- | --- | --- | --- | --- | --- |
| WT | None | 1018 ± 37 (8) | 720 ± 33 (7) | 0.71 | 3A |
| | AP4 | 1006 ± 42 (5) | 728 ± 56 (5) | 0.73 | NA |
| | AP4+PDA | 1044 ± 58 (5) | 878 ± 110 (5) | 0.84 | 3C, D; 6A-E; 7A, C |
| | Qcr | 896 ± 54 (3) | 977 ± 91 (4) | 1.09 | 4B, C; 5A-D |
| | Chq | 570 ± 53 (8) | 228 ± 44 (3) | 0.40 | 4B, C |
| | Pent | 990 ± 45 (5) | 295 ± 28 (5) | 0.30 | 4B, C |
| | VU590 | 770 ± 16 (6) | 264 ± 32 (6) | 0.34 | 4B, C |
| | Chq+Vu590 | 650 ± 32 (4) | 261 ± 32 (4) | 0.40 | 4B, C |
| | AP4+PDA+Qcr | 895 ± 50 (4) | 975 ± 134 (4) | 1.09 | NA |
| *Gnat2*<sup>-/-</sup> | None | 1009 ± 21 (3) | 814 ± 57 (3) | 0.81 | |
| | AP4+PDA | 1128 ± 20 (3) | 912 ± 48 (3) | 0.81 | 6A-E |
| | AP4+PDA+Qcr | 849 ± 67 (4) | 666 ± 150 (4) | 0.78 | NA |
| Avg |  | 1034 (24) | 790 (23) | 0.76 |  |

#### Resting level of K_o_ in the SRS

A general finding of the research supporting the OGSM hypothesis is that the “light-evoked decreased in (SRS) *K*_o_ can be mimicked *in vitro* in the isolated RPE-choroid preparation by decreasing apical *K*_o_ from 5 to 2 mM” (Gallemore et al., 1997). We thus adopted the value of 5 mM for resting SRS *K*_o_ in the model. It is notable that because 5 mM exceeds the level in mouse blood plasma (∼3.5 mM), the implication follows that ELM and RPE permeability barriers materially restrict K^+^ movement between the SRS and the choroidal and retinal vasculature, and that net K^+^ efflux from cells with plasma membranes in the SRS must exceed uptake (cf. Eq 4, first term on the right-hand side).

#### TER and TEP

A substantial body of results obtained from primary RPE cultures in Ussing chambers, and of isolated RPE cells constrain the model and its parameters. We prioritized parameter values extracted from reports of mouse RPE, but used results from other species when data from mouse was not available. The mouse RPE transepithelial resistance (TER) has been measured as *R*_RPE_ = 200 –400 Ω cm^2^ (Marmorstein et al., 2015; Fernandez-Godino et al., 2016), and given that the basal membrane resistance has been found to be substantially higher (1100 Ω cm^2^), TER was taken to be equivalent to *R*_Para_.

#### V_m, Ap_, V_m, Ba_ and TEP

The resting TEP of the RPE has been found to be 10 to 15 mV higher on the apical side, implying *V*_m, Ap_ to be 10-15 mV hyperpolarized relative to *V*_m, Ba_ (Eq 12; Fig. 2D). Measurements of the apical and basal membrane potentials have not to our knowledge been made in mouse RPE, but in frog *V*_m, Ba_ corresponds to *E*_K, SRS_ (Oakley, 1977)(Fig. 13), consistent with the results of (Shimura et al., 2001) that the conductance of Kir7.1 is the dominant apical conductance. *V*_m, Ap_ and *V*_m, Ba_ have been directly measured in frog RPE and bovine monolayers, and typical values are –80 mV and –70 mV, respectively.

#### E_K, SRS_, E_K, Ba_, E_Cl,Ba_

Based on the generally high permeability (low “reflection coefficient”) of capillaries to small ions and the further enhancement of the permeability by fenestrations (Michel, 2013), it can be reasonably assumed that the concentrations of the permeant ions Na^+^, K^+^, Cl^−^, Ca^2+^ in the ECS between the choriocapillaris and the basal RPE correspond to those of the blood plasma. Estimation of Nernst potentials require assignment of the cytoplasmic activities of the ions, and these have been taken from (Bialek and Miller, 1994). Similarly, we assumed that the dense band of retinal capillaries situated adjacent the ELM have sufficient permeability to peg the ECS concentration of the permeant at their plasma levels (**Fig. 2B**).

### Implementation of the model

*Rod ensemble model*. The portion of the model describing the molecular features and spatial distributions of rods membrane currents and their field potential are described in detail in (Peinado Allina, 2025b). In brief, the rod ensemble comprises 11 rods, whose inner and outer segments are identical but whose cell bodies are staggered over the outer nuclear layer. Rod outer segments express CNG channels and NCKX uniformly, and the inner segment expresses all the rod’s NKX, and K_v_2.1 and most of the HCN1 channels, and the remainder of the rod having a small unspecified K+ conductance and HCN1 channels (Fortenbach et al., 2021). For all simulations reported in this paper, it was assumed that 75% of the HCN1 channels reside in the inner segment membrane, with the remainder distributed uniformly over the rest of the cell according to membrane surface area. The electrical Equation of Continuity (EOC) for the dark steady-state was solved iteratively for computationally continuous axial spacing (0.5 μm). The rod ensemble was then discretized into 24 nodes spanning the photoreceptor layer and the dark steady-state re-established. (Peinado Allina, 2025b) to reduce the number of differential equations to be solved to compute the model light response. The delayed Gaussian model of the activation phase of phototransduction (Pugh and Lamb, 1993) for a 1 ms flash typically producing 3.5×10^5^ photoisomerizations rod^-1^ was used as the input to the model (cf (Peinado Allina, 2025b, Fig. 7), and it was assumed that the CNG-gated channels remained closed over the 3 s recording epoch (Supplementary Information, “2. Mouse rod responses in vivo to stimuli that generate saturating *c*-waves”). The rate equations governing NKX, Kv2.1 and HCN1 were modified from those used in (Peinado Allina, 2025b) to include explicit dependencies on SRS *K*_o_ (e.g.,. Eq 6).

#### Model of subretinal space

*K*_o._ Formulas for fluxes of K^+^ to/from the SRS are described in generically in Eqs 1–5, and the molecular details subsequently. For the model results presented here the SRS was assumed to be diffusionally equilibrated (“well-stirred”): thus, Eq 5 and subsequently provided ancillary relations defining the K+ fluxes comprised the rate equations for SRS *K*_o_. (In preliminary analyses K^+^ diffusion was included in the coding, but this inclusion complicated the achievement of the dark steady state of the whole model, requiring substantial baseline equilibration, and resulted in some instabilities, so was not extensively applied.)

#### RPE layer model

The dark steady-state of the RPE is characterized by Eqs 10 -24 (cf also the Appendix) and the assumption that in the dark SRS *K*_o_ = 5 mM. The electrical response of the RPE to changes in SRS *K*_o_ are described by the Eqs 25-26. (Additional information about the electrical response of the RPE layer to changes in SRS *K*_o_ are provide in the Supplementary Information, “3. Kinetic model of the RPE response to changes in SRS *K*_o_”).

#### Model of the Müller glial cell

The identification of the Slow PIII component of the ERG with currents of Müller cell Kir4.1 channels (Kofuji et al., 2000) and our own experiments showing that Slow PIII could be eliminated with quinacrine (**Fig. 5**), a Kir4.1-specific blocking drug (Marmolejo-Murillo et al., 2017a) led us to develop an electrical model of the Müller cell. The model rests on three factual principles: (1) Kir4.1 is by far the dominant conductance of the Müller cell (Kofuji et al., 2000); (2) Müller cells have extensive apical processes projecting that express Kir4.1 into the SRS across the ELM (**Fig S4.1**); (3) the multi-millivolt field potential generated by the rod dark current over the photoreceptor layer rises steeply over the axial extent of the Müller cell. A model based on these facts predicts that Müller cells generate a circulating current in the dark that is inward in the SRS region and outward elsewhere in the cell, with both inward and outward current flowing through Kir4.1 channels (Supplementary Material, “4. Müller glial cell circulating current in the dark and light”). Because the rod field potential rapidly collapses and reverses direction in response strong light (Peinado Allina, 2025b), and SRS *K*_o_ subsequently declines from its resting level, the Müller glial cell is also predicted to have a light-driven field potential response with at least two distinct components, one ephatic and a second tracking the SRS decline in SRS *K*_o_ (cf. Supplementary Information, “Electrophysiological Model of the Mouse Müller Cell and the Extraction of Slow PIII”).

**Figure 5.**
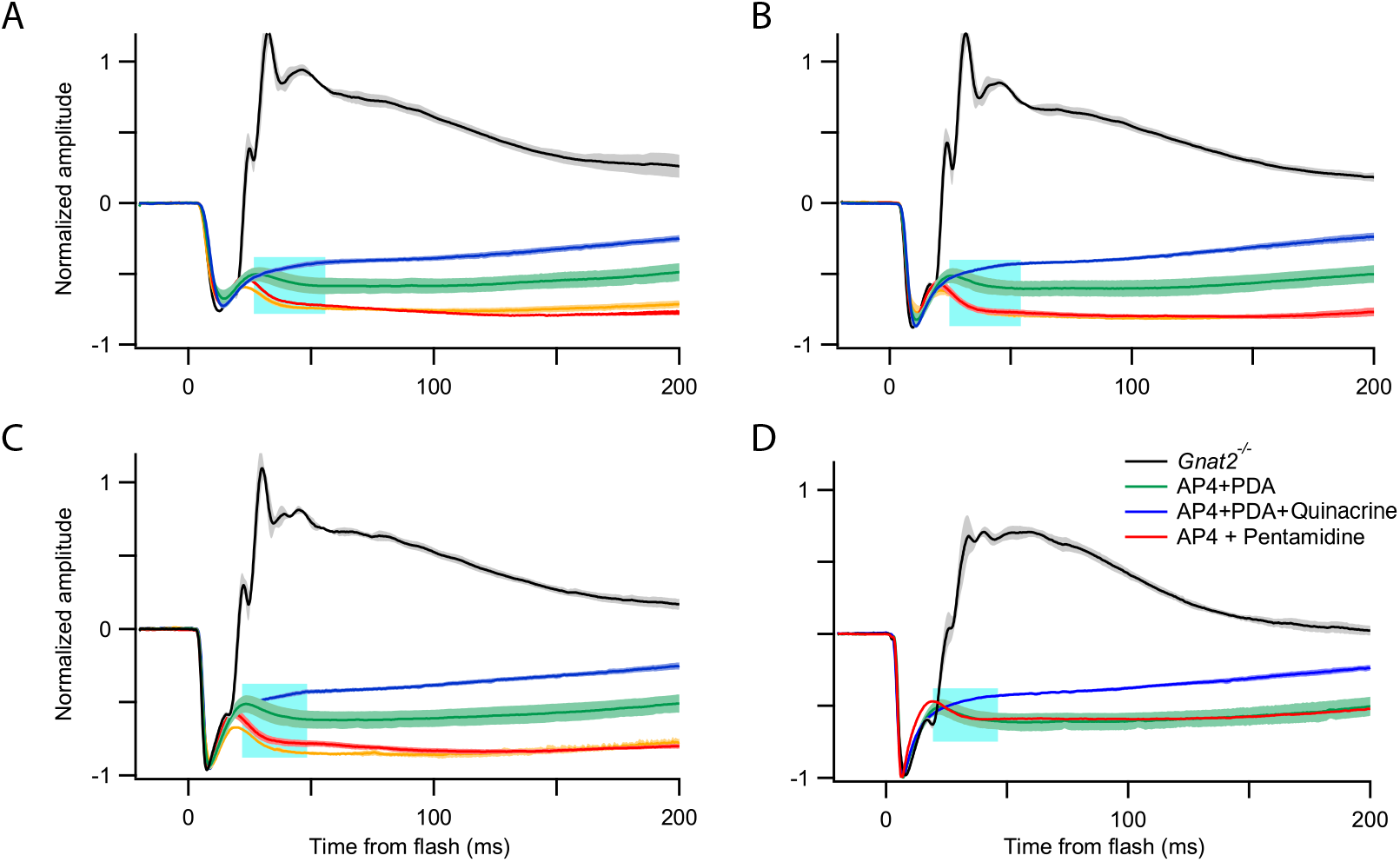
Quinacrine blocks the Slow PIII component of the mouse ERG. Each panel shows the average ERGs of *Gnat2*^−/−^ mice exposed to a 1 ms flash, and intravitreally injected with the drugs indicated in the legend of panel D and WT controls. Flash energy densities (in photons μm^-2^) were as follows: 9.3×10^3^ (A), 3.7×10^4^ (B), 1.5×10^5^ (C), 6×10^5^ (D). The negative-going “Slow PIII” region of the figures has been framed by a cyan rectangle. For all flash strengths, the “Slow PIII” component is absent after intravitreal injection of quinacrine, an established pore-blocker of Kir4.1 (Table 1). (Error bands around traces are SEM’s, with n β 3.) The positive-sloping segment of the ERG measured in the presence of AP4+PDA+Quinacrine (blue trace) is the beginning of the observable *c*-wave (cf. Supplementary Information, “4. Müller glial cell circulating current in the dark and light”).

#### Units

Physical units and symbol conventions follow those in (Peinado Allina, 2025a; Peinado Allina, 2025b) (cf **Table 2**), with distinctions needed between the physical units in the “continuous” modeling of the dark steady-state, and the “discretized” modeling of both the dark steady state and the dynamical response to light stimulation. To accommodate Müller cells’ dark steady state, the model was extended from the OPL to the GCL, including 16 additional nodes in the discretized model, for a total of 40 nodes spanning the retina. The discretized model subdivides the retina into 40 nodes, 6 in the rod outer segment region, 5 in the rod inner segment region, and 13 for the ONL and OPL, with 16 for the remainder of the retina (cf. Fig. 2, symbols), with each node cumulating the membrane current over a defined mutually exclusive axial extent, so that rod contributions to nodal currents had units of A rod^-1^. Nodal current was converted to current density (A cm^-2^) by multiplying by the rod density, ρ_rods_ = 3.5 × 10^7^ rod cm^-2^. Müller cell currents’ contributions to the nodal currents were scaled by 1/22, based on the mouse Müller cell density of 16,000 cm^-2^ measured by (Chao et al., 1997). For the 11 most distal nodes, which are assigned to the SRS, K^+^ current density was converted to flux density (mols cm^-2^) by dividing by Faraday’s constant, with a specialized treatment used to extract the unidirectional K^+^ current from HCN1 channels, which are permeable to both K^+^ and Na^+^ (Supplementary Information, “1. Calculating K^+^ flux through mouse rod HCN1 channels”). Membrane currents of RPE cells were naturally expressed as current densities (A cm^-2^), and RPE apical K^+^ flux was included in “node #1”, which also included the rod outer segment tips (“ROST”). Fluxes densities of K^+^ across the ELM (node 11) and RPE (node 1) barriers were computed as described by Eqs 2, 3. Finally, the rate of change of *K*_o, SRS_, the concentration of K^+^ in the “well-stirred” SRS, was obtained by summing the K^+^ flux densities of the SRS nodes and dividing by *V*_SRS_, the SRS volume density (L cm^-2^) (Eq 5).

#### Coding

The model was coded in Matlab™. The 564 differential equations comprising the dynamic model were numerically integrated with the routines “ode23” or “ode23s” with a fixed time step of 100 μs on the UC Davis Franklin High Performance “Hippo” server; a run covering 3 s of model time typically took about 8 hrs of Hippo time.

## RESULTS

### K^+^ fluxes in the mouse subretinal space in the dark steady-state are dominated by rod ionic mechanisms

As a requisite first step in predicting light-driven changes in SRS *K*_o_, a model must predict dark steady states of the currents and ion fluxes of rods, RPE and Müller cells, and simultaneously yield a zero net SRS K^+^flux. The resting value of SRS *K*_o_ was set to 5 mM based on the studies cited in the Introduction, while the level of *K*_o_ outside the SRS was taken to be that of blood plasma, 3.5 mM (**Fig. 2B**). The latter assumption was rationalized by the abundance and proximity of the choroidal and outer plexiform layer vasculatures, and the negligible reflection coefficient of capillaries for K^+^ (Michel, 2013). The distribution of *K*_o_ provided the basis for assigning *E*_K_ in the three cell types both in and outside the SRS (**Fig. 2D**). Müller cells have apical processes that extend into the inner segment region of the SRS and have a resting potential dominated by highly expressed Kir4.1 channels (Kofuji et al., 2000), and are thereby expected to have a resting potential very near *E*_K_. This expectation was met by the model prediction to within less than 3 mV over the length of the Müller cell (**Fig. 2D**; cf. Supplementary Information, “4. Electrophysiology Model of the Mouse Müller Cell and the Extraction of Slow PIII”.) However, the electrochemical driving force for K^+^ reverses sign for the portion of the Müller cell membrane facing the SRS versus that in the rest of the retina, predicting a resting circulating current that is inward in the apical processes of the SRS and outward elsewhere. While the corresponding extracellular current density of the Müller cells is small relative that of the rods, it is predicted to generate an extracellular axial potential (**Fig. 2E**, magenta symbols), one that is sensitive to both the light-dependent decrease in the rod field potential and to the decline in SRS *K*_o_ that occurs during strong light exposures.

With the distribution of *K*_o_ in darkness specified, the electrical and ionic steady-states of the three cell types and the K^+^ fluxes to/from the SRS were found by an iterative search process that also balanced the net K^+^ flux to/from the SRS (Theory, “Implementation of the Model”). The rod inner segment NKX and K_v_2.1 channels, which combine to generate the bulk (∼85%) of the steady-state outward electrical current that balances the inward current of the rod outer segment (Fortenbach et al., 2021; Peinado Allina, 2025b), have opposite-signed K^+^ fluxes that dominate and nearly neutralize the total SRS K^+^ flux (**Fig. 2F**).

### Blocking rod synapses and genetic elimination of cone signaling leaves the murine c-wave intact

The flash-activated mouse *c*-wave is a light-dependent, corneal-positive transient whose maximal magnitude in the untreated eye is comparable to those of the *a*-wave and *b*-waves, but which only appears after subsidence of the *b*-wave in the untreated retina (**Fig. 3B**). The flash-activated *c*-wave has the same or a slightly increased saturated positive peak-amplitude when synaptic transmission from photoreceptors is pharmacologically blocked and cone responses genetically eliminated (*Gnat2*^−/−^) (**Fig. 3C, D; Table 3**). In addition to its light-dependence (**Fig. 3D**) and its persistence in when the *b*-wave is pharmacologically eliminated, a feature of the mouse flash-activated *c*-wave linking it to rod responses is the correlation between the duration of the rod photoresponses measured *in vivo* under identical conditions with the paired-flash method (Peinado Allina et al., 2017) and the time when the *c*-wave reaches peak amplitude and begins recovery: thus, the longer that rods remain in saturation, the slower the *c*-wave recovery (compare same colored traces in **Fig. 3D** and **3F**, respectively) (cf. Supplementary Information, “2. Recovery of mouse rod responses *in vivo* to stimuli that generate **c**-waves”). This correlation is qualitatively consistent with the OGSM hypothesis, which predicts that while CNG channels remain closed by ongoing phototransduction, NKX activity will continue to pump SRS K^+^ into the rod IS, lowering SRS *K*_o_.

### Pharmacological blockers of Kir7.1 but not of Kir 4.1 channels greatly reduce the murine c-wave

Molecular specification of the OGSM hypotheses requires identification of the RPE K^+^ channels that respond to photoreceptor-driven decline in SRS *K*_o_. Kir7.1 channels are strongly expressed in the mouse RPE apical membrane, but nowhere else in the retina (**Fig. 4A**) (Shimura et al., 2001; Yang et al., 2003). To test the hypothesis that Kir7.1 channels play a major role in the generation of the mouse *c*-wave, selective Kir channel inhibitors were introduced intravitreally with and without synaptic blockers, and their effects on the principal components of the ERG measured. The Kir7.1-selective inhibitor VU590 (**Table 1**) reduced the *c*-wave on average to 34% of its normal value (**Fig. 4C**; **Table 3**). Chloroquine and pentamidine also severely reduced the *c*-wave, though evidence for their ability to block Kir7.1 channels in patch-clamp data is lacking.

A complication of the use of chloroquine and VU590 to test the role of Kir7.1 channels in *c*-wave generation is that these drugs also reduced the *a*-wave (by 45% and 25%, respectively), suggesting that the rod dark current was reduced. However, pentamidine reduced the *c*-wave to 30% of its normal amplitude while negligibly (4%) reducing the *a*-wave (**Fig. 4B, C; Table 3**). This effect of pentamidine is consistent with hypothesis that pharmacological reduction of the *c*-wave generator current for an hour need not materially alter the resting rod circulating current, but also suggests that chloroquine and VU590 have a direct or indirect effect on the rod circulating current – e.g., by inhibiting Kv2.1 channels or NKX activity, or by interfering with a role of Müller cell Kir4.1 in setting the resting SRS *K*_o_ .

### The “slow-PIII” component of the ERG is blocked by quinacrine

In the presence of the glutamatergic synaptic blockers AP4 and PDA, immediately after the fast *a-*wave relaxation attributable to extracellular flow of rod capacitive current (Robson and Frishman, 2014; Peinado Allina, 2025b, Fig. 3C) there is a corneal-negative ERG transient classically identified as “Slow PIII” (this nomenclature arose from historical identification of the corneal negative transient now called the *a*-wave as “Fast PIII”). The slow-PIII component of the ERG has long been attributed to Müller cells, and given that Kir4.1 channels contribute more than 90% of the resting conductance of mouse Müller cells (Kofuji et al., 2000), Slow PIII should be reduced or eliminated by pharmacological block of Kir4.1: this is indeed the case for the Kir4.1-specific drug quinacrine (Marmolejo-Murillo et al., 2017a) (**Fig. 5; Table 3**). Notably, quinacrine has no effect on the *c*-wave (**Fig. 4C**), and is associated with only a small (13%) reduction in the *a*-wave (**Table 3**). This latter reduction may be ascribable to the loss of a fast ephatic component of the Müller cell response, which is expected to have a waveform very similar to the early rod *a*-wave (cf. Supplementary Material, “4. Electrophysiological Model of the Mouse Müller cell and the Extraction of Slow PIII.”)

In summary, intravitreal injections of photoreceptor synaptic blockers and Kir-selective blockers support the hypothesis that Kir7.1 expressed in the apical RPE and Kir4.1 expressed in Müller cells (**Fig.4A**) respectively play essential roles in the generation of the *c*-wave and slow PIII components, respectively, of the mouse ERG.

### Quantitative features of the mouse c-wave recorded in the presence of synaptic blockers

The mouse *c*-wave is readily observable in ERG recordings made in the presence of the photoreceptor synaptic blockers, AP4 and PDA, which eliminate post-synaptic components of the ERG (**Fig. 3C, D**; **Fig. 6**). These results not only affirm that physiological events post-synaptic to photoreceptors play a negligible role in the generation of the *c*-wave, they also allow key quantitative features of the *c*-wave to be extracted more readily and fully. These features include: (1) the maximum amplitude (**Fig. 6B, D**), (2) the maximum rate of rise (**Fig. 6C, D**); (3) the time delay to the appearance of the *c*-wave (**Fig. 6D**); (4) the dependence of the time to criterion recovery on flash strength (**Fig. 6B, E**).

**Figure 6.**
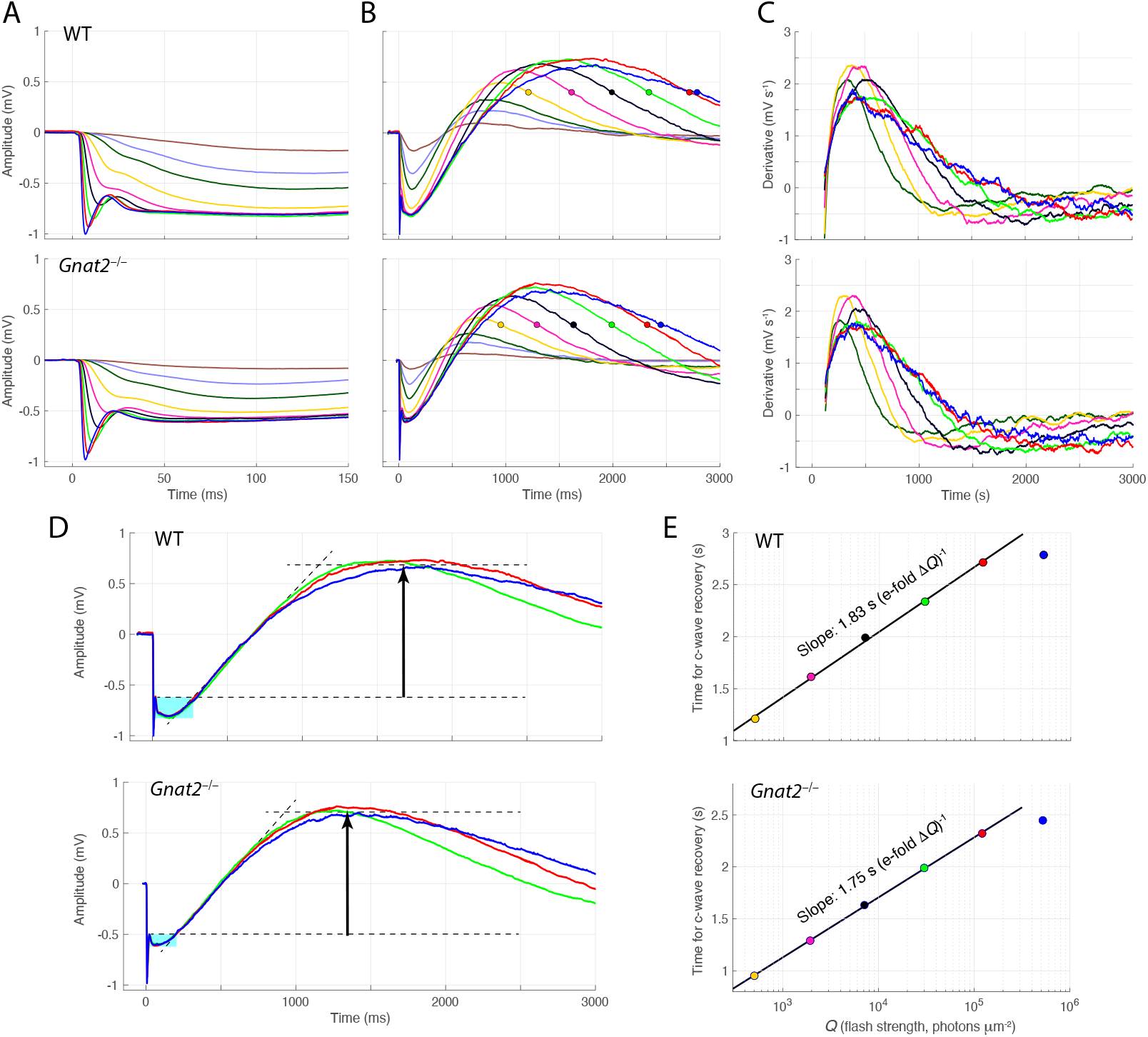
Quantitative features of flash-activated ERG *c*-wave response families obtained in the presence of synaptic blockers (AP4, PDA). **A, B**. Averaged ERG response families of WT and *Gnat2*^−/−^ mice to flashes delivering 6.2 to 500,000 photons μm^-2^ shown on 150 ms (A) and 3000 ms (B) time scales. In the initial ∼ 400 ms the negative-going *a*-wave dominates, but the positive-going *c*-wave dominates the records for all flash strengths after about 500 ms. **C**. Derivatives of the traces in panel B for the six most intense flash strengths. **D**. Saturation of the *c*-wave. ERGs measured in response to the 3 most intense flashes are plotted on an expanded scale: until approximately 700 ms the traces have identical rising trajectories, and differ minimally up to about 1300 ms. **E**. “Pepperberg plot” (Pepperberg et al., 1992; Lyubarsky et al., 1996; Lyubarsky and Pugh, 1996; Hetling and Pepperberg, 1999) of the *c*-wave recoveries: the times when each trace recovers to a criterion level (colored circles in B) are shown as a function of the flash strength on a semilogarithmic plot: the slopes (obtained by least-squares fitting) provide an estimate of a “dominant time constant” governing the timing of the *c*-wave recoveries in this intensity regime. The topmost point in each plot deviates from the fitted lines and is a manifestation of saturation: thus, a 4.3-fold increase in flash strength produces negligible differences in the rising phase and in the timing of the early recovery (compare red and blue traces in panels B and D).

#### Saturation of the c-wave

The initial 500 ms of the ERG light response family measured in the presence of glutamatergic synaptic blockers appears at first glance to resemble rod photocurrent responses (**Fig. 6B**). However, the response recovery phases contravene the expected monophasic character of rod responses (**Fig. 3F**), even for the weakest flash (6.2 photons μm^-2^; brown trace). These overshoots get increasingly large with increasing flash strength, and unequivocably transition into the *c*-wave. The *c*-waves recorded in response to the three strongest flashes saturate with respect to flash strength in three distinct ways: latency of appearance, velocity, and amplitude. Rod responses to such stimuli do not exhibit these properties, but rather continue to increase in duration with flash strength over a much larger range (cf. **Fig. 3F**), a behavior caused by the increasing quantity of non-deactivated photoactivated rhodopsin.

#### Maximum amplitude

In the presence of the synaptic blockers, the mouse *c*-wave has a maximum amplitude from baseline of 0.6 to 0.7 mV. However, the saturated *c*-wave rises during the *a*-wave “plateau”, i.e., after the fast relaxation of the *a*-wave from its negative minimum to a negative plateau potential of – 0.6 mV (dashed lines in **Fig. 6D**). Consequently, the saturated *c*-wave has an amplitude under these experimental conditions of 1.2 to 1.3 mV.

#### Maximum velocity

Unlike the rod-driven *a*-wave, the *c*-wave has similar rising phase kinetics over a greater than 100-fold range in stimulus strength (**Fig. 6B**). The somewhat higher initial velocity of *c*-waves (**Fig. 6B, C**) generated by flashes that do not produce maximal *a*-wave plateaux may seem anomalous, but is attributable to *c*-wave generation co-occurring with recovery of the rod response: i.e., the field potentials of two distinct positive-going sources with are combining. For the strongest flashes, however, the *a*-wave generator can be presumed not to undergo any recovery in the 3 s epoch and so will not distort the *c*-wave rise (cf Supplementary Information, “2. Mouse rod responses *in vivo* to stimuli that generate saturating *c*-waves”). Thus, the maximum rate of rise of the pure *c*-wave can be taken to be the value obtained in the responses to the three most intense flash strengths, ∼1.7 mV s^-1^ (**Fig. 6C**, green, blue and red traces).

#### Light-dependence of c-wave recoveries

Insight into the duration of rod responses during the epoch (500 to 3000 ms) of the corneal-positive phase of the *c*-wave is provided by examination of the recovery (late, negative-going) phase of the *c*-waves, which are seen to be displaced along the time axis in a shape-invariant manner (**Fig. 6B**). Photoreceptors are the only retinal cells previously known produce such shifting behavior in response to increasing intensity flashes, and it is reasonable to conclude that the lateral shifts in *c*-wave recovery on the time axis are caused by underlying shifts in rod response recovery. This conclusion is strengthened by “Pepperberg” plots of the recovery times as a function of the logarithm of the flash strength (**Fig. 6E**). The slopes of the plots provide estimates of the time constant of an underlying an approximately exponential decline, which we attribute to decay of rhodopsin (“R*”) catalytic activity during this epoch. The extracted time constant, 1.8 s for both WT and *Gnat2*^−/−^ mice, is close to values previously obtained from recordings of single WT rods (Krispel et al., 2006)(1.5 s), and from two paired-flash studies of rod *a*-waves (Lyubarsky and Pugh, 1996; Peinado Allina et al., 2017) (1.6 s and 2.0 s respectively) for flashes producing between 3000 to 10^5^ photoisomerizations per rod (see Supplementary Information, “3. Mouse rod responses *in vivo* to stimuli that produce saturating *c*-waves”). Accordingly, it can be concluded that the timing of the recovery phase of the *c*-waves in response to flashes delivering up to ∼10^5^ photons μm^-2^ is determined by the recoveries of the rod responses.

#### Latency in appearance of the c-wave

Another distinctive feature of the synaptically isolated, flash-activated mouse *c*-wave is its delay or latency of appearance: thus, extrapolating the maximal slope of the saturated *c*-wave to the level of the *a*-wave plateau one finds an intersection at 200 to 250 ms that serves to define the latency (**Fig. 6D**). A clue to the nature of the delay is provided by the appearance of the negative-going “Slow PIII” transient in this time window (cyan-rectangles).

### Prediction of the pharmacologically isolated, saturated mouse c-wave

The saturated mouse *c*-wave can be approximately predicted by the model developed in this investigation to embody the OGSM hypothesis (**Fig. 7**). The model component field potentials are generated by three cell types with plasma membranes facing the SRS: (i) rods; (ii) RPE cells; (iii) Müller cells. The empirical saturated responses are first presented (**Fig7A**), and then the theoretical and summed source photopotentials (**Fig 7B**). The magnitude of each source is governed by two distinct types of factors, intrinsic and extrinsic. Intrinsic factors include molecular mechanisms (channels and transporters) that determine the local currents at the cells’ membranes, while extrinsic factors include the conductivity distribution and layer conductances of the eye and extraocular tissues (Peinado Allina, 2025a). In the following we describe the ways in which the model simulations illuminate the roles of these factors.

**Figure 7.**
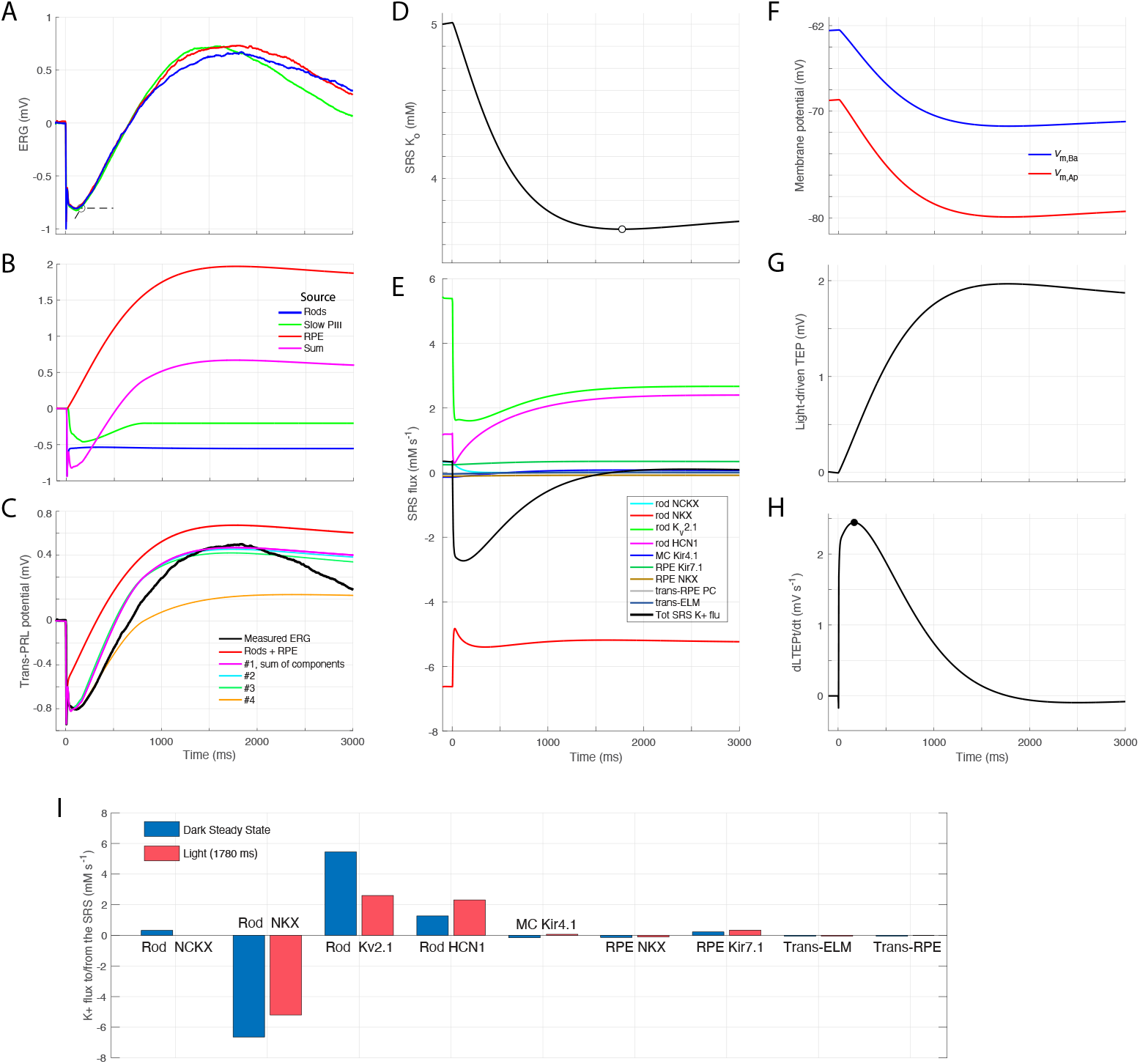
Prediction of the saturated mouse *c*-wave measured in the presence of synaptic blockers. **A**. ERGs obtained in response to the three most intense flashes of Fig. 6D. **B**. Sources of the three cells of the model (rods, RPE, Müller cells) presented scaled for the corneal ERG. The rod source is the trans-receptor layer response for a strongly saturating flash producing of 3.5×10^5^ photoisomerizations rod^-1^; mouse rods exhibit no recovery in 3 s from such a stimulus, reflected in the rod source’s (blue curve) negligible change after the initial fast capacitive relaxation (Robson and Frishman, 2014). The Müller cell source was obtained as the “Slow PIII” component (Supplementary Information, **Fig. S4.3**). The RPE source is described in panels F, G. The magenta curve is the sum of the three components. **C**. Prediction of the *c*-wave. The black (data) curve is the average of the red and blue traces from A. The red curve presents the sum of the rod and RPE sources from panel B, and serves to emphasize that the sum of these two components rises with negligible delay on this time scale. However, when the Müller cell source is included in the sum, the predicted trace (magenta) is brought into much closer agreement with the data trace, as the latter shifts the red trace downward. Three other examples (#2 – #4) of model calculations employing different parameters are shown. **D**. Time course of SRS *K*_o_ light-driven change predicted by the model; the open symbol indicates the minimum, which occurs at 1377 ms. **E**. Components of K^+^ flux to/from the SRS predicted by the model expressed in molar units. **F**. RPE apical (*V*_m,AP_) and basal (*V*_m,Ba_) membrane potentials predicted by the model. The “driving force” for these changes is the decline in SRS *K*_o_ (panel D), which serves as the input to the RPE membrane potential rate equations, Eqs 25, 26 (cf Supplementary Information, “3. Kinetic model of the RPE response to changes in SRS *K*_o_”.) **G**. The RPE trans-epithelial potential (TEP) obtained as the difference of the curves in F; the TEP serves as the RPE source for the prediction (panel B) when appropriately scaled. **H**. Derivative of the TEP: this shows that the model maximum velocity occurs early, and provides as a benchmark for comparison with *c*-wave data. **I**. Bar-chart comparison of SRS K^+^ flux components in the dark steady state (blue; cf. Fig. 2F) with the K^+^ fluxes from the same components during the “light-driven” pseudo-steady-state defined by the time of the minimum of SRS *K*_o_ (panel D, open symbol).

#### Rod ensemble source

The ionic mechanisms of the model rod ensemble and the consequent field potential response to saturating flash were presented in (Peinado Allina, 2025b), and are here extended to a 3s epoch (**Fig. 7B**, blue curve). The scaling between the field potential “at the source” and the response measured at the cornea was examined in (Peinado Allina, 2025b), and will be taken up again in the Discussion. The link between the rod photoresponse and the *c*-wave is the rod-driven decline in SRS *K*_o_ (**Fig. 7D**) (Oakley and Green, 1976; Oakley, 1977; Steinberg et al., 1980) which shifts RPE apical *E*_K_ (cf. Supplementary Information, “Kinetic model of the RPE response to changes in SRS *K*_o_”).

#### The decline in SRS K_o_ caused mainly by reduction in rod K_v_2.1 current

The predicted light-stimulated change in SRS *K*_o_ (**Fig. 7D**) is governed by 9 distinct components of K^+^ flux (**Fig. 7E**; Eqs 5–9). As in the dark state **(Fig. 1F**) throughout the period of decline of *K*_o_ from its resting level the rod NKX component has the largest magnitude, and remains relatively constant throughout the *c*-wave after a rapid initial flux decrease caused by rod hyperpolarization. The second-most prominent dark component, K^+^ efflux through K_v_2.1 channels, decreases almost 3-fold as rod hyperpolarization closes these channels: thus, the model reveals that the rapid decline in SRS *K*_o_ is caused mainly by the decline in K_v_2.1 current (**7C**, green curve) –in fact, due to its dependencies on membrane potential and *K*_o_ (Stanley et al., 2015; Fortenbach et al., 2021), the magnitude of the K^+^ uptake from the SRS into rods by the NKX declines by 20% between the dark state and 1 second after the stimulus, slowing the rate of *K*_o_ decline. A histographic snapshot of the component K^+^ fluxes at the bottom of the decline (∼ 1380 ms) further serves to emphasize the key role of the decline of K_v_2.1 current (**Fig. 7I**).

#### The RPE source tracks the rod-driven decline in SRS K_o_

The RPE transepithelial potential (TEP) (**Fig. 7E, G)** tracks the change in SRS *K*_o_. The mechanistic link between *K*_o_ and the TEP is the negative shifting of the apical RPE K^+^ Nernst potential: this causes the Kir7.1 I/V curve to shift correspondingly, and thereby increasing outward current through Kir7.1 channels, which in effect act as a K^+^ electrode for the SRS (Supplementary Information, “3. Kinetic model of RPE response to changes in SRS *K*_o_”, **Fig. S3.1**).

#### Müller glial cells’ “light response” (Slow PIII) underlies the apparent delay in the rise of the c-wave

The rapid (∼ 35 ms) hyperpolarization-driven closure of 70% of the inner segment K_v_2.1 channels and continued NKX activity is predicted to cause SRS *K*_o_ to undergo a linear decline thereafter, and produces a serious conflict between theory and data (**Fig. 7C**, red vs. black curves). The appearance around the time of the *c*-wave delay of “Slow PIII”, indubitably attributable to Müller cells (Kofuji et al., 2000) -- suggests that the Müller cell electrical response might underlie the *c*-wave delay. While a completely satisfactory model of the Müller cell’s “light response” has not yet been achieved, the “Slow PIII” response (**Fig. 7C**, green curve) was extracted from subtraction of the synaptically isolated ERGs measured in the presence and absence of the Kir4.1 blocker quinacrine (Supplementary Information, “4. Müller glial cell circulating current in the dark and light”). Summing this Slow PIII component the model’s rod (blue) and RPE (red) curves provides a reasonable account of most of the delay (**Fig. 7C**, magenta curve).

#### Key extrinsic factors

Key extrinsic factors affecting the component magnitudes include the “internal series resistance” of each source (e.g., *R*_Para_ the paracellular resistance across the RPE). The values adopted are consistent published with mouse data (**Table 2**). One notably important extrinsic factor affecting the kinetics and magnitudes of all components of SRS *K*_o_ change is the volume of the SRS (cf. Eq 5, *V*_SRS_): with all other parameters held constant, the time at which *K*_o_ reached its minimum was found to be linearly (affine) dependent on *V*_SRS_ over the latter’s plausible range.

#### A surprising turnabout

The model predicts SRS *K*_o_ to begin to recover to baseline around 1380 ms after the saturating stimulus: this recovery occurs while the rod source remains constant (**Fig. 7B**, blue curve), and does not arise from deactivation of rod phototransduction (cf Supplementary Information, “2. Mouse rod responses *in vivo* to stimuli that generate saturating *c-*waves”). The two principal causes of this turnaround are the increases in K_v_2.1 and HCN1 currents (**Fig. 7E**, green and magenta curves), whose reversal potentials are changed by the decline is SRS *K*_o_. These channels thus constitute a feedback mechanism for driving the recovery of *K*_o_. Smaller contributions to the turnaround are made by the decreasing Nernst potentials of the Müller glial cell Kir4.1 and RPE Kir7.1 channels. In addition, when *K*_o_ declines below the blood plasma level both trans-ELM and trans-RPE extracellular K^+^ influxes into the SRS increase (**Fig. 7G**), as these depend on the *K*_o_ differences across the barriers (Eqs 2, 3).

## DISCUSSION

### The Subretinal Space (SRS) is a bounded extracellular space (ECS) that encompasses most of the photoreceptors’ circulating currents

The SRS is a contiguous volume of ECS that bathes the densely packed layer of photoreceptor outer and inner segments (**Fig. 2**)(Steinberg, 1985b). The SRS is bounded on its proximal side by the external limiting membrane (ELM), and on its distal side by the retinal pigmented epithelium (RPE) layer, long recognized as a major blood-retinal barrier of the posterior eye. Rod photoreceptors in the dark-adapted state generate a massive extracellular current, ∼600 μA cm^-2^ at the junction between the current’s inner segment sources (K_v_2.1, NKX) and outer segment sinks (CNG channels, NCKX) (Hagins et al., 1970; Peinado Allina, 2025b), Figs. 5, S1). Because the fractional volume of the SRS is only a few percent of the total inner-outer segment layer volume, and K^+^ fluxes through the NKX and K_v_2.1 channels are large in molar units (**Fig. 1F**, ∼ 6 mM s^-1^), light responses that completely close the CNG channels, hyperpolarize the cell and thereby close K_v_2.1 channels can cause SRS *K*_o_ to undergo physiologically material changes (**Fig. 7**).

The highly fenestrated choriocapillaris and abundant choroidal vasculature serve as potent regulators of the ionic composition of the ECS bathing the basal RPE. Similarly, immediately proximal to the ELM is a layer of retinal vasculature, which like the choriocapillaris can be expected to have a low reflection coefficient for permeant ions (Michel, 2013), “pegging” the ECS ionic composition immediately adjacent close to that of the blood plasma. Thus, the two vasculature layers bracketing the SRS appear located so as to maintain ion concentrations in the immediately adjacent ECS close to those of the blood plasma, and for the ELM also to isolate the neurons of the inner retina from SRS *K*_o_ fluctuations. These ideas are embodied in the hypothesized resting distribution of *K*_o_ (**Fig. 2B**).

### Molecular mechanisms of the Oakley-Green -Steinberg-Miller (OGSM) theory of the ERG c-wave

A classic body of physiological research supports the hypothesis that the source of the *c*-wave of the ERG is a transient transepithelial potential (TEP) of the retinal pigment epithelium (RPE). For example, light-driven changes in *K*_o_ in the subretinal space (SRS) measured with a K^+^-selective electrode perfectly track the simultaneously measured RPE TEP (Oakley and Green, 1976; Oakley, 1977; Steinberg et al., 1980). Based on an accrued wealth of physiological investigations of isolated mammalian RPE cells and primary RPE cultures e.g., (Bialek and Miller, 1994; Shimura et al., 2001), and the histochemically measured distributions of mouse rod ionic mechanisms (Fortenbach et al., 2021; Peinado Allina, 2025b), we developed a molecularly specified, biophysical model of the mouse *c*-wave, and compared its predictions with the ERG measured in the presence of synaptic blockers (**Fig. 7**). Though yet imperfect, the predictions by the model provides strong support the OGSM hypothesis (**Fig. 7**), and provide insight into the contributions of different cellular and molecular mechanisms that determine SRS *K*_o_.

### Evaluating the 3-source model of the synaptically blocked ERG in response to intense flashes

The model comprises three electrical sources whose field potentials sum to predict the synaptically blocked ERG over a 3 second epoch that includes the saturated *c*-wave (**Fig 6D**): rods, Müller cells and the RPE (**Fig. 7B**).

#### Rod source

The rod source (**Fig. 7B**, blue curve) is essentially constant over the *c*-wave epoch. Its negative transretinal potential is a consequence of current flowing outward through NKX and K_v_2.1 channels of the inner segment that sinks inward through HCN1 channels: while ∼ 80% of the total 9 pA per rod outward current returns through HCN1 channels in the inner segment (and thus does not contribute materially to a transretinal potential), the residual 20% re-enters through HCN1 channels distributed over the outer nuclear layer (Peinado Allina, 2025b, Fig. 5C, F, I). Thus, abundant HCN1 sinks made available by these channels’ eponymous hyperpolarization consequent to the closure of the outer segment CNG-gated channels causes a net extracellular current that flows toward the anterior retina (i.e., positively in the standard retinal axial coordinate), creating a negative field potential across most of the photoreceptor layer. The rod source potential at the cornea, ∼ – 0.6 mV was obtained by scaling the trans-retinal rod source by the same factor as was used in prediction of the *a*-wave (Peinado Allina, 2025b) (**Fig. 7**).

#### RPE source

The RPE source (**Fig. 7B**, red curve) is a transepithelial potential (TEP) across the paracellular resistance of the ∼ 5 μm thick RPE monolayer (**Fig. 1**). (Eqs 25-26; cf Supplementary Information, “3. Kinetic model of the RPE response to changes in SRS *K*_o_”). The highly expressed Kir7.1 channels located in the RPE apical membrane (Shimura et al., 2001; Yang et al., 2003; Shahi et al., 2017; Cornejo et al., 2018) are the molecular transducers of the TEP, acting as sensors for the light-driven decline in SRS *K*_o_. Such decline predictably shifts the K^+^ Nernst potential at the RPE apical membrane, increasing outward current of Kir7.1 channels, which passes through the paracellular resistance *en route* to chloride conductance sinks, Bestrophin and CFTR, in the basal membrane (Adijanto et al., 2009) (**Fig 1E**). As the apical and basal membrane potentials hyperpolarize in response to the increased outward apical current (**Fig. 7F**), the TEP increases by ∼ 2 mV (**Fig. 7G**). The parameters of the RPE model are close to those measured in experiments on mammalian RPE cells (**Table 2**), and the source-to-cornea scaling factor close to that expected from the 4-shell volume current model of the mouse eye (Peinado Allina, 2025a).

#### Müller cell source

The “Slow PIII” component of the ERG (**Fig. 7B**, green curve; cf also **Fig. 3C, Fig. 4, Fig. 5D**) is firmly attributed to extracellular current carried by Müller cell Kir4.1 channels: specifically, Slow PIII is absent in *Kir4*.*1*^−/−^ mice (Kofuji et al., 2000), and was eliminated by the Kir4.1-specific blocking agent, quinacrine (**Fig. 5; Table 1**)(Marmolejo-Murillo et al., 2017a), which negligibly affects the *c*-wave (**Table 3**). It may seem surprising that the Müller cell could show a response to light given that ∼90% of its conductance is determined by Kir4.1 channels (Kofuji et al., 2000), and that it receives no trans-synaptic signals from photoreceptors. However, Müller cells are “downstream” from photoreceptors in two distinct ways, as exemplified by analysis of the dark steady state. Firstly, the field potential of the rod circulating current (**Fig. 2E**, black symbols) rises several mV from the distal to the proximal portion of the Müller cell, with the consequence that the membrane potential of the proximal portion is hyperpolarized with respect to the distal portion in the dark adapted state (**Fig. 2D**, blue symbols and curve). Secondly, the apical processes of the Müller cells extend into the SRS where *K*_o_ is elevated relative to its concentration in the ECS of the more proximal retina (**Fig. 2C**). The consequence of these facts is that Müller cells have a circulating current in the dark state of the retina that is inward in the SRS and outward elsewhere: thus, *I*_Kir4.1_(*x*) = *g*_Kir4.1_(*x*) [*V*_m_(*x*) – *E*_K_(*x*)] is negative in the SRS and positive in the outer nuclear layer (**Fig. 2D**). Moreover, these ideas lead to the expectation that Müller cells should exhibit two components of light-response: (1) an “ephatic” component arising from the “collapse” of the rod field potential; (2) a “K^+^-sensing” component in which the photoreceptor-driven decline of SRS *K*_o_ alters *E*_K_ of the Müller cells’ apical processes, and it is reasonable to hypothesize that (1) and (2) are the mechanistic basis of “Slow PIII” (Kofuji et al., 2000). Unfortunately, while we were able to predict the dark circulating current of the Müller cells and the consequent field potential “ab initio” (**Fig. 2E**, magenta symbols; cf, Supplementary Information, “4. Müller glial cell circulating current in the dark and light”), we have been unable to-date to successfully derive their light response, and so chose instead to use as the Müller cell source in the 3-source model an empirical source obtained by subtraction of ERGs obtained in the presence of AP4, PDA and quinacrine from ERGs measured with AP4 and PDA only (**Fig. 7B**, green curve; **Fig. S4.3**).

#### Conclusions about and Perspective on the 3-source ERG model

We conclude overall that the summed potentials of the three cellular sources (**Fig. 7B**, magenta curve) provide a reasonable if yet imperfect “3-source” prediction of the synaptically blocked, saturated mouse ERG *c*-wave (magenta vs black curves in **Fig. 7C**), a prediction consistent with the OGSM hypothesis. One aspect of this conclusion is that the negative-going Müller cell source is the likely explanation of the delay in the rise of the *c*-wave, subtracting from the RPE TEP positive-going source that tracks the decline in SRS *K*_o_ which begins shortly after the stimulus. A second interesting aspect is that all the SRS K^+^ flux components have “built-in” negative feedbacks that act to slow and even reverse the decline in SRS *K*_o_ (**Fig. 7E**). These feedbacks act either through negative shifts in their local *E*_K_ (rod K_v_2.1 channels, Müller cell Kir4.1 and RPE Kir7.1 channels; Eqs 11, 13), through photoreceptor hyperpolarization activation (HCN1 channels), through the dependence of NKX activity on *K*_o_ (rod and RPE NKX; Eq 10), or via trans-barrier concentration differences of *K*_o_ (ELM; RPE paracellular path; Eqs 2, 3 ).

The present model is naturally extendable to include features that should improve its description of the synaptically blocked, saturated mouse ERG *c*-wave, including the histochemically measured inhomogeneous distributions of K*v*2.1 and NKX in the rod inner segments ((Peinado Allina, 2025b), combined with diffusion of K^+^ in the SRS. The dominant sources and sinks of SRS K^+^ flux reside in the inner segment, and histochemical profiles reveal both to be concentrated closer to the ELM. As a consequence, including diffusion of K^+^ in the SRS can be expected to increase the delay with which the NKX-driven decline in SRS *K*_o_ reaches the Kir7.1 sensors in the apical RPE membrane and further shift the predicted response forward in time. Implementing a successful dynamical model of the Müller cell response – to be tested against the pharmacologically dissected Slow PIII (**Fig. S4.3**) -- should also contribute to improving the model. In particular, an ephatic component of the Müller cell source caused by the negative-going early phase of the rod *a*-wave can be expected to closely track the rod source field potential, as the mouse Müller cell has a membrane time constant of ∼ 1 ms (25 MΩ × 50 pS). If the 3-source model can be made to closely describe the synaptically blocked, saturated mouse ERG *c*-wave, it should be straightforward to extend it to include the recovery phase of the rod current, using a Pepperberg constant of ∼ 2 s (**Fig. 6E, Fig. S2**) as an estimate of the deactivation time constant of photoactivated rhodopsin.

### A role for rod HCN1 channels in SRS K_o_ balance

The principal extant hypothesis for the function of rod HCN1 channels is that the depolarization consequent to their activation serves to restore the membrane potential at the synapse towards its normal operating range. The localization of ∼75% of the rod’s HCN1 channels in the inner segment suggests another important role, viz. that HCN1 in the inner segment (i.e., facing the SRS) plays an key role as a feedback mechanism for SRS *K*_o_ homeostasis, acting to restore *K*_o_ towards its resting level when activated by hyperpolarization (**Fig. 7E**, magenta trace).

### Dissecting with drugs the corneal ERG and its reconstitution with a molecular/biophysical model

This investigation contributes to a body of evidence showing that drugs injected intravitreally into the mouse eye can reach their molecular targets in the posterior retina while producing negligible effects on a widely used benchmark of rod function, the ERG photovoltage *(a*-wave). Intraocular injection of the glutamatergic synaptic blockers AP4 and PDA, previously used successfully in dissecting ERG components (Kang Derwent et al., 2007; Robson and Frishman, 2014), establishes that the *c*-wave retains its principal features in the absence of synaptic transmission (**Figs 3, 7**), and further reveals the full magnitude and kinetics of the saturated *c*-wave when the rod response is saturated (**Fig. 7D**). Overall, the results and modeling illustrate the translational era’s program of integrating the vast wealth of molecular-scale ion channel and transporter physiology and meso-scale tissue characterization into “*in silico*” models of neuronal function *in vivo*. The model presented here also provides a foundation for further understanding retinal function *in vivo*, for non-invasive testing of animal models of disease and for the evaluation of therapeutic interventions.

## APPENDIX. Derivation of the Steady-state of the Retinal Pigment Epithelium

The RPE model cell comprises the following components.

### Variables

*a* = *J*_NKX_ (electrogenic current through NKX)

*b* = *J*_K_ (all K^+^ channel current)

*c* = *J*_Cl_ (all Cl^−^ channel current)

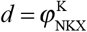 (K^+^ influx through NKX)

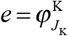 (K^+^ efflux through *J*_K_ )

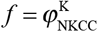 (K^+^ influx through NKCC)

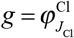 (Cl^−^ efflux through *J*_Cl_ )

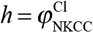 (Cl^−^ influx through NKCC)

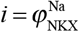 (Na^+^ efflux through NKX)

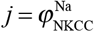 (Na^+^ influx through NKCC)

### Equations

0 = *a* + *b*+ *c*(1) (charge balance)

0 = *d* + *e* + *f*(2) (K^+^ flux balance)

0 = *g* + *h*(3) (Cl^−^ flux balance)

0 = *i* + *j*(4) (Na^+^ flux balance)

*d* = −2*a* / *F*(5) (NKX stoichiometry, current→ K^+^influx)*e* = *b* / *F*

(6) (*J*_K_ current→ K^+^efflux)*f* = *h* / 2

(7) (NKCC stoichiometry)*g* = −*c* / *F*

(8) (*J*_Cl_ current→Cl^−^ efflux)*i* = 3*a* / *F*

(9) (NKX stoichiometry, current→ Na^+^efflux)*j* = *h* / 2

(10) (NKCC stoichiometry)

### Solution steps

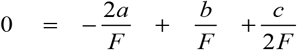 (11) (Substitution from Eqs 3, 5-7 in Eq 2)

0 = − 3*a* − *c* / 2 (12) (multiply Eq 11 by F; subtract Eq 1)

*c* = − 6*a* Solution from previous *c* in terms of *a*

### *Complete solution in terms of J*_NKX_

*a* = *J*_NKX_ can be selected based on measurements and constraints

*c* = − 6*a*

*b* = 5*a* (interesting that *J*_K_ must be 5 × *J*_NKX_ )

*d* = −2*a* / *F*

*e* = *b* / *F* = 5*a* / *F*

*f* = − *c* / 2*F* = − 3*a* / *F*

*g* = −*c* / *F* = 6*a* / *F*

*h* = *c* / 2*F* = − 6*a* / *F*

*i* = 3*a* / *F*

*j* = *h* / 2 = − 3*a* / *F*

### Check solution against the conservation equations

*a* + *b* + *c* = *a* + 5*a* − 6*a* = 0

*d* + *e* + *f* = (−2*a* + 5*a* − 3*a*) / *F* = 0

*g* + *h* = (6*a* − 6*a*) / *F* = 0

*i* + *j* = (3*a* − 3*a*) / *F* = 0

This analysis is based purely on electrical and mass conservation, and transporter stoichiometry, and does not involve the apical/basal location of the seven molecular components (**Fig. 1E**).

#### 3. Steady-state “model” of the RPE TEP: constraints from biophysics and measurements

Some constraints taken into consideration in setting model parameters are as follows:

i. measurements of NKX flux or electrogenic current (*J*_NKX_)(Shimura et al., 1999);
ii. measurements of the TER (*R*_Para_) (multiple studies);
iii. measurements of the apical/basal distribution of K+ current *α* = *J*_Kir7.1, rest_ / *J*_K, tot_ ; (Bialek and Miller, 1994) characterize the apical vs. basal distribution of *J*_K_.
iv. measurements of the resting TEP: *TEP* = *R*_Para_ ( *J*_NKX_ + *J*_Kir7.1, rest_ ) = *R*_Para_ *J*NKX (1+ 5*α* ) ;
v. measurements of apical or basal membrane potentials (one or the other, given TEP)

#### 4. Dependencies on V_m_

These come into play in the SRS dynamics – e.g., the Kir7.1 I/V curve is important for the prediction of the apical outward current triggered by SRS *K*_o_ decline (and the consequent *E*_K,SRS_ shift) (cf Table 2 and Supplementary Information, “. The dark SS sets the absolute scales for all the ionic mechanisms at the respective apical and basolateral *V*_m_’s: e.g., *g*_Cl_ comes out close to that measured by Bialek & Miller (1974).

FIGURES for Peinado *et al*., “Rod, RPE and Muller cell contributions to light-driven changes in SRS *K*_o_”

## SUPPLEMENTARY INFORMATION

### 1. Calculating K^+^ <lux through mouse rod HCN1 channels

Mammalian rod HCN1 channels have a reversal potential of ∼ –30 mV (Demontis et al., 1999; Demontis et al., 2002; Demontis et al., 2009), revealing them to be permeable to both Na^+^ and K^+^. Since they are highly expressed in the rod inner segment (Peinado Allina, 2025b), a model of *K*_o, SRS_ needs to include K^+^ flux through HCN1 channels. There is (to our knowledge) no generally accepted model of the HCN1 K^+^ permeability for the membrane potential range over which mouse rods function (–32 mV to –75 mV). To meet the need for estimating HCN1 K^+^ flux, we used the Goldman-Hodgkin-Katz current (GHKC) relation as a computational tool (**Fig. S1**), nonetheless for its limitations (Lewis, 1979).

### 2. Recovery of mouse rod responses in vivo to stimuli that generate saturating c-waves

The studies whose data are reproduced in Fig. S2 show that for flashes estimated to produce between 3000 and 10,000 photoisomerizations rod^-1^, recoveries from CNG channel current-saturating flashes undergo a change of law. Thus, below this cutoff, recovery times can be characterized as being rate-limited by a “dominant” time constant of 200 to 250 ms, while above the cutoff the recovery times are governed by a time constant of 1500 to 2000 ms. In the present study, *c*-wave recovery times exhibited Pepperberg behavior with a dominant time constant of ∼ 1800 ms (**Fig. 6B**), consistent with the idea that the timing of the *c*-wave recoveries (**Fig. 6E**) is determined by rod CNG-gated channel current coming out of saturation. Saturated *c*-wave responses were recorded in response to stimuli estimated to deliver 10^5^ to 10^6^ photons μm^-2^ (**Fig. 6F**): given an end-on mouse rod collecting area *in situ* of 0.5 to 1 μm^2^ (Lyubarsky et al., 2004), the rod responses underlying the saturated *c*-waves are expected to exhibit no recovery in 3 seconds. Thus, in addition to establishing comparability of the rod “Pepp2” and *c*-wave dominant recovery time constants, the data in Fig. S2 support the assumption made for the model of the saturated *c*-wave that after the few milliseconds during which CNG channels close, rod CNG-channel current remains in saturation throughout a 3 s recording epoch (**Fig. 7A, C**).

### 3. Kinetic model of the RPE response to changes in SRS K_o_

The OGSM RPE model (Eqs 10–26) posits that the during the flash-activated ERG Kir7.1 channels of the apical membrane act as K^+^ sensors. This SI section provides a description of the RPE model’s intrinsic kinetics (see **Fig. 1** for an anatomical description of the mouse RPE cell layer, along with schematic of the molecular components and their disposition in the apical and basal membrane (Adijanto et al., 2009).

We extracted the current/voltage (I/V) relation of Kir7.1 channels measured in primary bovine RPE cultures as the barium-sensitive current (Shimura et al., 2001); these I/V data were fitted with a Boltzmann relation to provide an analytical description (**Fig. S3.1A**, black curve). The I/V curve was then shifted to have a zero crossing at *E*_K_ = –72.4 mV (Bialek and Miller, 1994), with *K*_o,SRS_ = 5 mM, *K*_i_ = 75 mM and *V*_m,Ap_ = –69 mV (Bialek and Miller, 1994), and conductance scaled so that the current density of Kir7.1 channels was 17.5 μA cm^-2^ (corresponding to an NKX current density of 2.5 μA cm^-2^ measured by (Shimura et al., 1999) (cf the Appendix). It can then be readily calculated how the Kir7.1 I/V curve would be shifted at the instant that *K*_o,SRS_ “jumps” to a lower concentration by shifting the K^+^ Nernst potential, as seen here for *K*_o,SRS_ jumps from 5.0 mM to lower values (**Fig. S3.1B)**. These shifted I/V curves show how increases of outward Kir7.1 result from dynamical decline in *K*_o,SRS_ (**Fig. S3.1C)**.

Kir7.1 channels and the NXK are the dominant ionic mechanisms of the apical RPE and generate outward current that cannot be balanced by inward current from apical sources. Rather, the outward apical current “sinks” (i.e., is balanced) as inward current through the chloride-sensitive Best and CFTR channels of the basal membrane (**Fig. 1E**).

### 4. Electrophysiological Model of the Mouse Müller Cell and the Extraction of Slow PIII

Features of mouse retinal Müller cells pertinent to their contributions to the ERG are as follows. Müller cells span most of the retina, from their apical processes in the inner segment layer (the proximal SRS) to the vitreal surface (**Fig. 4**; **Fig S4.1**), and have a spatial density in mouse of 16,000 mm^-2^ (Chao et al., 1997), 1:22 relative to the rod density assumed in the model (Peinado Allina, 2025a). The axo-dendritic process of the mouse Müller cell has been estimated to have an average total resistance of 32 MΩ, about 2.2 MΩ μm^-1^ (Dmitriev et al., 2021). Kir4.1 channels are ubiquitously expressed in Müller cells (**Fig. 4A**), and are responsible for more than 90% of the resting conductance: in whole-cell recordings the average input resistance of the wild-type mouse Müller cell is 25 ± 3 MΩ, increasing ∼ 22-fold to 311 MΩ in Müller cells of *Kir4*.*1*^−/−^ mice measured at the WT resting potential, −85 mV (Kofuji et al., 2000).

We used high-resolution confocal immunohistochemistry (IHC) with a Kir4.1 antibody to derive axial retinal expression density profiles of Kir4.1 (**Fig. S4.1**); the integral of the profile multiplied by the reciprocal of the Kir4.1 input resistance, 40 nS, provides the axial distribution of the Kir4.1 conductance (nS μm^-1^).

Other membrane proteins that play important roles in the Müller cell’s electrical and ionic homeostasis include the electrogenic Na^+^: HCO_3_^−^ co-transporter NBC1 (mouse protein Nbc1; gene, *Slc4a7*), the NKX anti-porter, and the excitatory amino-acid co-/counter-transporter EAAT1 (mouse: Eaat1; gene *Slc1a3*) (**Table S4.1**). The rationale for including Nbc1is that the distal Müller cell very strongly expresses carbonic anhydrase, which traps as HCO_3_^−^ much of the CO_2_ generated by photoreceptor aerobic respiration. The cell then transports as HCO_3_^−^ to the retinal vasculature, where Nbc1 delivers it (after reconversion to CO_2_) to hemoglobin in red blood cells (Newman, 1994; Chiang et al., 2022). The efflux of Na^+^ via the rod inner segment NKX that balances the influx through CNG-channels and outer segment NCKX for a rod with a dark current of 20 pA rod^-1^ (Fortenbach et al., 2021) requires hydrolysis of 0.071 fmol s^-1^ rod^-1^ of ATP. Assuming the ATP is generated by aerobic metabolism – 32 ATP’s generated per glucose molecule and 6 CO_2_ molecules produced per glucose molecule – maintenance of the dark current generates 0.013 fmol s^-1^ rod^-1^ of CO_2_. Assuming 100% of the rod-generated CO_2_ is trapped as HCO_3_ ^−^ in the Müller cells (1 per 22 rods) and exported by the Nbc1 co-transporter with a stoichiometry 2HCO_3_^−^ : 1Na^+^, the Nbc1 electrogenic current would be –14.2 pA per Müller cell in the dark adapted retina. For the Müller cell to maintain Na^+^ homeostasis in the presence of the loss via Nbc1, an influx is required, and this appears to be provided by Eaat1 (Glast1). In addition to Kir4.1 and the three aforementioned electrogenic tranporters, the MC also expresses three non-electrogenic transporters, KCC (K^+^: Cl^−^ co-transporter), AE3 (Cl^−^: HCO_3_^−^ exchanger), MCT (monocarboxylate transporter), and the electrogenic V-ATPase. The stoichiometries and transport direction of all the transporters are summarized in **Table S4.1**.

The axial distributions of Nbc1, NKX and Eaat1 in the Müller cell were assigned according to the cell’s membrane surface area distribution and the transporter function. Thus, for example, Nbc1 was assumed to be expressed primarily in the vascularized outer plexiform, inner plexiform and ganglion cell layers (**Fig. S4.2**, OPL, IPL, GCL). Give the axial distributions of Kir4.1, Nbc1 NK and Eaat1 (and the assumed distribution of *K*_o_ in the ECS – **Fig. 2**), it was possible to determine axial distributions of the Müller ionic currents and membrane potential that satisfied the Equation of Continuity (“charge conservation”), and the corresponding ECS current density and field potential (**Fig. S4.2**).

In addition to satisfying the EOC, the Müller cell must satisfy ion-species homeostasis (ISH) to be in a steady-state in the dark. The problem of satisfying both the EOC and ISH was solved in the main text for the RPE (cf Appendix), but is more challenging for the Müller cell in part due to the effects of the axial distributions of *K*_o_ and *V*_m_ on the Kir4.1 current, and in part due to lack of the same level of information about the different transporters’ velocities (cycles per second). If however, the velocities of the transporters are taken as a vector of 9 unknowns, 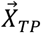, the problem can be reduced to a series of 8 linear equations in the 9 unknown velocities, with 7 of the equations expressing the ISH for the ions and the 8^th^ the EOC. We tried to find solutions to these equations subject to various additional constraints (e.g., capture as the carbonic anhydrase product HCO_3_^−^ of all or part of the CO_2_ generated by rod aerobic respiration), and while no perfect solution for the 7 ISH equations has yet been found, it is notable that the EOC can be achieved readily by the Kir4.1 current distribution in all reasonable cases of the transporter activities. Given the membrane currents found to satisfy the EOC, the membrane potential (**Fig. 4.2E)**, membrane currents (**Fig. 4.2F)** and field potential (**Fig. 4.2H**, red curve) of the Müller cell in the dark adapted state were readily computed.

*Extraction of Slow PIII from ERG recordings*. We were unable to predict the Müller cell’s “light response” by including appropriate differential equations in the master set of equations governing the rod ensemble, SRS *K*_o_ and the RPE. This failure likely arose from the “stiffness” of Müller cell rate equations, in particular the relatively low capacitance in the SRS region, and can probably be remedied by a more suitable choice of variables -- e.g., *U*_MC_(*x, t*) = *c*_m,MC_(*x*) *V*_m,MC_ (*x, t*), rather than *V*_m,MC_ (*x, t*). The possibility that the Müller cell-generated “Slow PIII” component of the ERG (Kofuji et al., 2000) might be responsible for the delay in the rise of the *c*-wave (**Fig. 5D**) led us to extract it from ERGs made in the presence of the synaptic blockers AP4 and PDA with and without quinacrine, which blocks Kir4.1 channels (**Fig. S4.3**) …

*Expectation of and evidence for a Müller cell ephatic response that is a scaled, inverted mimic of the rod field potential*. The dark current (**Fig. S4.2G)** and steady-state field potential (Fig. 2E, of the Müller cell (**Fig. S4.2**) rests on three foundations: (1) the ubiquitous expression of Kir4.1 channels (**Fig. 4.1**) and their dominance of the Müller cell’s conductance (Kofuji et al., 2000); (2) the elevated concentration of K^+^ in the SRS (**Fig. 1B**), which cause *E*_K_ across Müller cell processes in the SRS to be higher than elsewhere **(Fig. 1D, Fig. S4.2E**); (3) the relatively rapid increase in the field potential of the rods over the most distal Müller cell, which causes the distal *V*_m_ to be elevated relative to that in proximal portion by dint of the *V*_o_ (**Fig. S4.2E)**. These features also predict that the light response of rods will generate two distinct corneal-negative changes in the Müller cells’ field potential: (1) an ephatic response arising from the rod-driven change in *V*_o_; (2) a slower change arising from the decline of SRS *K*_o_. We believe that these two distinct components are seen in the Slow PIII response extracted from the synaptically blocked ERGs obtained with and without quinacrine (**Fig. S4.3**). However, we have as yet been unable to generate predictions of the Müller cell field potential with the model, and so assignment of the apparent components of pharmacologically isolated Slow PIII (and predicting the response in the first 20 ms after intense rod stimulation) remains an ongoing challenge. Some evidence for the ephatic component of the Müller cell response comes from paired comparison of the *a*-wave amplitudes measured in the presence of AP4+PDA, but with and without the Kir4.1-current blocking agent quinacrine (Qcr^−^ vs Qcr^+^) (**Table 3**): for WT the difference in amplitude is 149 μV (*t* = 1.99, *p =* 0.045, 7 df, 1-tailed test); for the *Gnat*^−/−^ mice, the difference is 279 μV (*t* = 3.44, *p =* 0.009, 5 df, 1-tailed test. For the pooled results of the two genotypes, the average *a*-wave amplitude difference is 204 μV (*t* = 3.68, *p =* 0.001, 14 df, 1-tailed test). This difference was neglected in extracting Slow PIII (**Fig. S4.3A**): rather the traces were scaled so that the *a*-waves matched. Inclusion of the absolute scales implies that the Müller cell ephatic response has essentially the same waveform as the rod-driven *a*-wave over the initial 12 ms, as the waveforms closely match for the Qcr^−^ vs Qcr^+^ conditions. It bears note in this context that the model Müller cell conductance was assigned to Kir4.1 and fixed at 40 nS (Kofuji et al., 2000), and that the steady-state analysis invariably predicted a source potential at the cornea of +200 to +300 μV (**Fig. 1E, Fig. S4.2H**), regardless of reasonable choices for the cell’s electrogenic transporters: in achieving EOC neutrality, the model cell reacted to increased inward electrogenic current by increasing the outward Kir4.1 current. Thus, it can be anticipated that a successful Müller cell dynamic model will not only generate an ephatic response with the form of the initial negative-going portion of the rod *a*-wave, but that the magnitude of this response at the cornea will be 200 to 300 μV. Finally, these ideas may contribute to the understanding of the overall physiological function of the Müller cell: the dominance of the conductance by Kir4.1 appears to give the cell the ability to react efficiently to varying transporter demands, such as one might expect given that carbonic anhydrase in the distal cell likely traps a large fraction of the CO_2_ generated by photoreceptor aerobic respiration.

**Figure S1.**
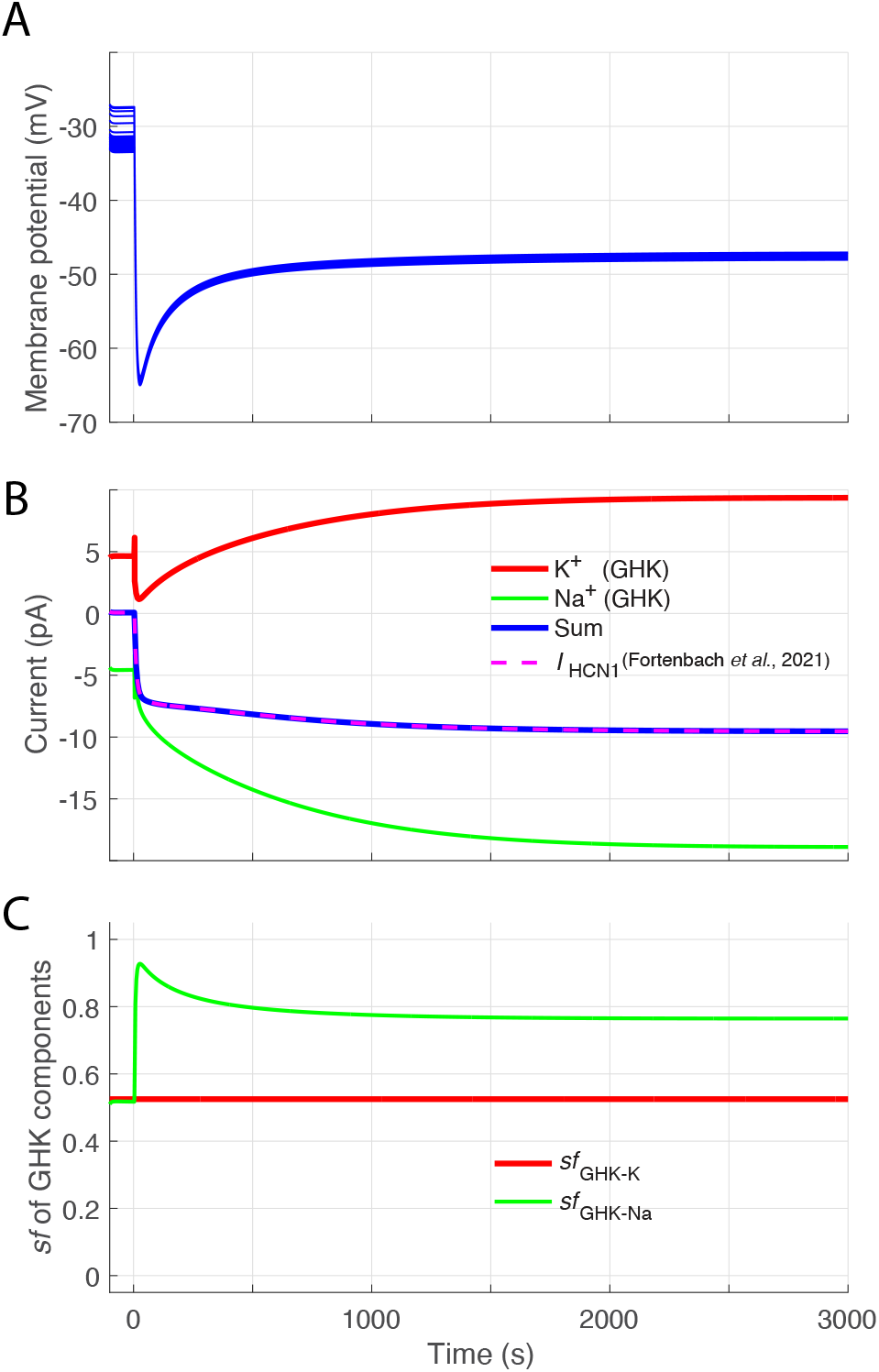
Calculation of K^+^ fluxes through rod HCN1 channels. **A**. Model-generated rod hyperpolarizing photovoltage (PV) response to a flash that suppresses the CNG channel current for at least 3 s. The PV traces are the average over the 11-rod ensemble for 24 axial locations spanning the photoreceptor layer (the dark adapted rod is not isopotential, and dispersion in traces for *t* < 0 occurs in the outer segment locations (cf. (Peinado Allina, 2025b)). **B**. HCN1 current of the average rod driven by the PV response in A, as calculated with 2-state Boltzmann model of HCN1 (dashed magenta curve) (Fortenbach et al., 2021) The red and green curves represent unidirectional K^+^ and Na^+^ currents computed by setting the GHK current formula (Eq 9) equal to the prediction of the 2-state Boltzmann model at each time point (each *V*_m_); the K^+^ component of the GHK current was scaled by a factor (*sf*_GHK-K_ = 0.525), and the blue curve is the sum of the red and green curves: thus, the unidirectional currents were made consistent with the prediction of the 2-state Boltzmann model. **C**. Plot of the scaling factors for the two GHK component currents: the K^+^ current component was scaled by the constant *sf*_GHK-K_, while the Na^+^ component was dynamically modified so the sum of the two components equaled that predicted by the 2-state Boltzmann. In fitting the model to *c*-wave data, *sf*_HCN1_ was varied between 0.525 and 0.7. (For *sf*_GHK-K_ < 0.525 component summation failed.)

**Figure S2.**
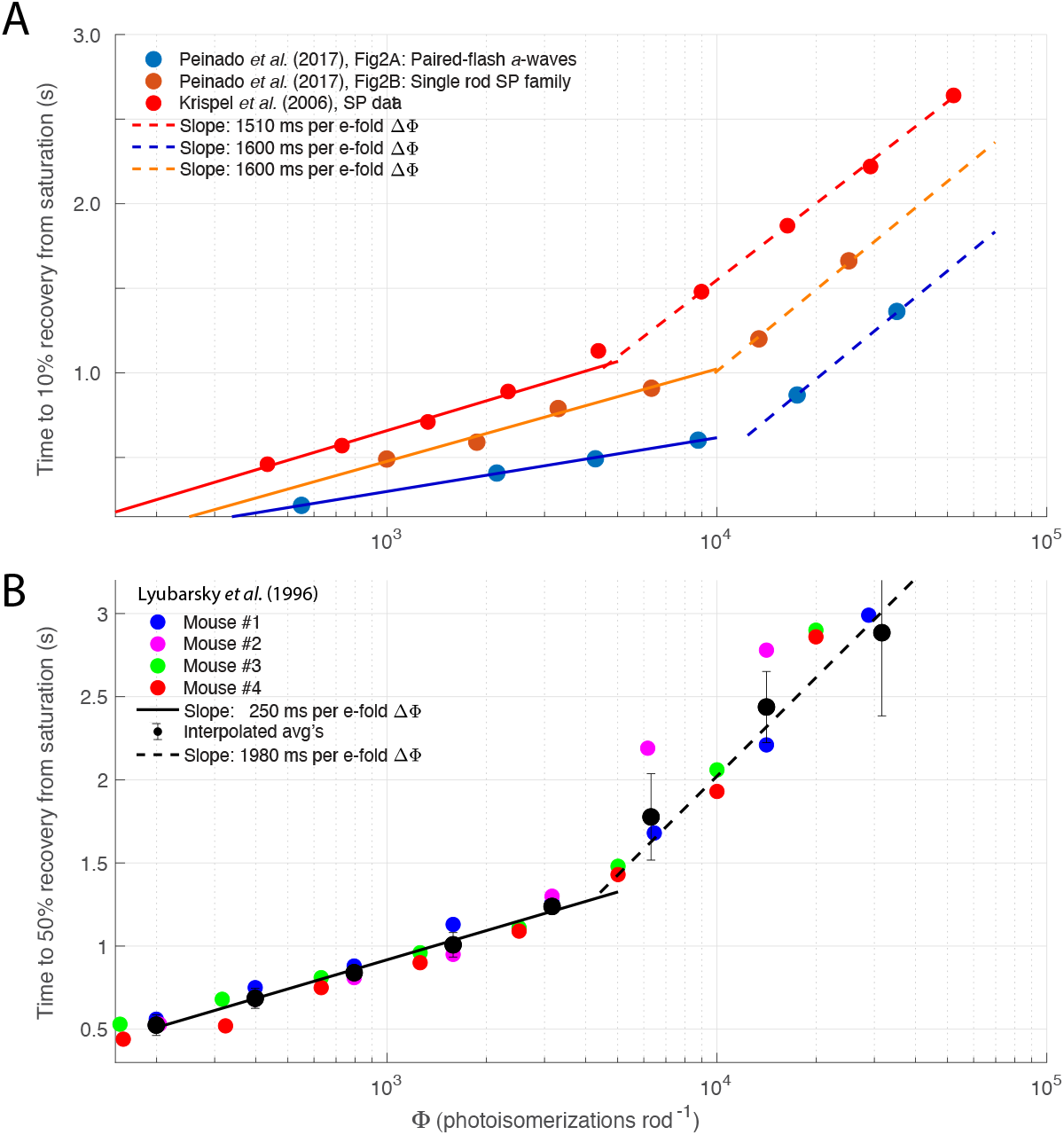
Analysis of data from several studies showing that rod response recovery is greatly slowed in the “Pepperberg 2” stimulus energy regime. **A**. Pepperberg plots (time to criterion recovery of rod dark current plotted vs. logarithm of stimulus energy) from 3 studies of the recovery of mouse rod photo-currents, including one with the paired-flash ERG *a*-wave (Peinado Allina et al., 2017), and two with suction pipette recordings from rod outer segments (Krispel et al., 2006) (Peinado Allina et al., 2017). **B**. Pepperberg plot of data from *in vivo* paired-flash ERG *a*-waves (Lyubarsky and Pugh, 1996).

**Figure S3.1.**
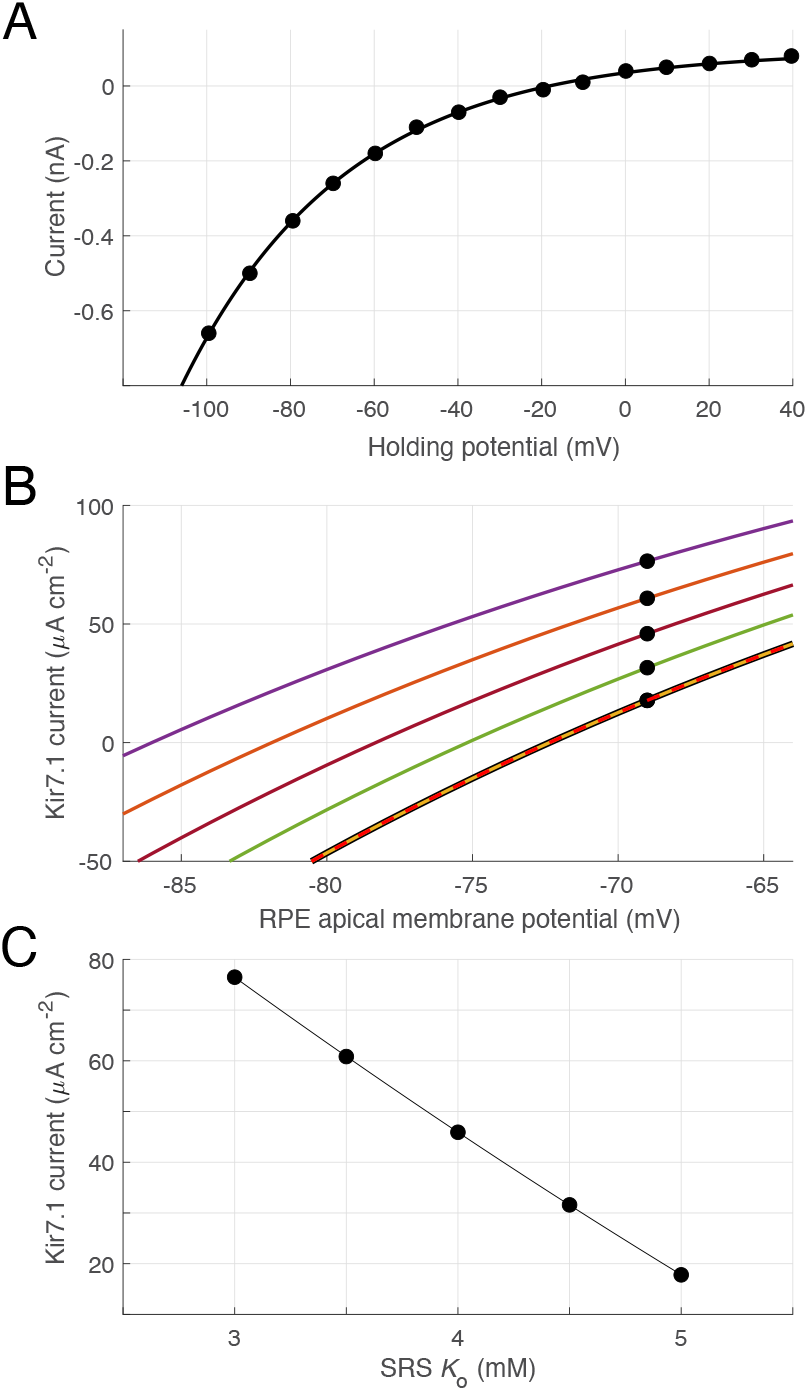
Characterization of RPE Kir7.1 channel current. **A**. Current/voltage relation of the barium-sensitive current of bovine RPE cells measured by (Shimura et al., 2001) (Fig 10A; 140 mM K^+^) (filled symbols); the abscissa gives the holding potential (*V*_h_), and the ordinate the measured clamp current. The data were fitted with a Boltzmann relation (smooth curve), *I*_Kir7.1_(*V*_h_) = *I*_Kir7.1_(*V*_h0_){1–exp[-(*V*_h_-*V*_h0_)/*s*_Kir7.1_]}, with, *V*_h0_ = – 60 mV, *I*_Kir7.1_(*V*_h0_) = – 0.18 nA and *s*_Kir7.1_ = 40 mV. **B**. The Boltzmann function of A was shifted to have its zero-crossing (K^+^ Nernst potential) at – 72.4 mV, and scaled to have a current density at the RPE apical membrane resting potential – 69 mV (Bialek and Miller, 1994) of 17.5 μA cm^-2^ (thickened red curve). The other curves plot the I/V curve shifted laterally for K^+^ Nernst potentials for SRS *K*_o_ between 5.0 mM and 3.0 mM in 0.5 mM steps (the zero-crossings are at the Nernst potentials). **C**. RPE Kir7.1 apical membrane current density for different SRS *K*_o_ values (the black symbols replot the symbols whose ordinate values are seen in B). The smooth curve is a quadratic regression function obtained with the Matlab “polyfit” function: the dominant, linear component has a slope of -39 mV per mM increase in *K*_o_. In the model, selection of the RPE parameter *J*_NKX, RPE, rest_ (the magnitude of the resting Kir7.1 current dictated the initial Kir7.1 I/V curve prior to light stimulation, in conjunction with the other RPE-specific parameters (Appendix).

**Figure S3.2.**
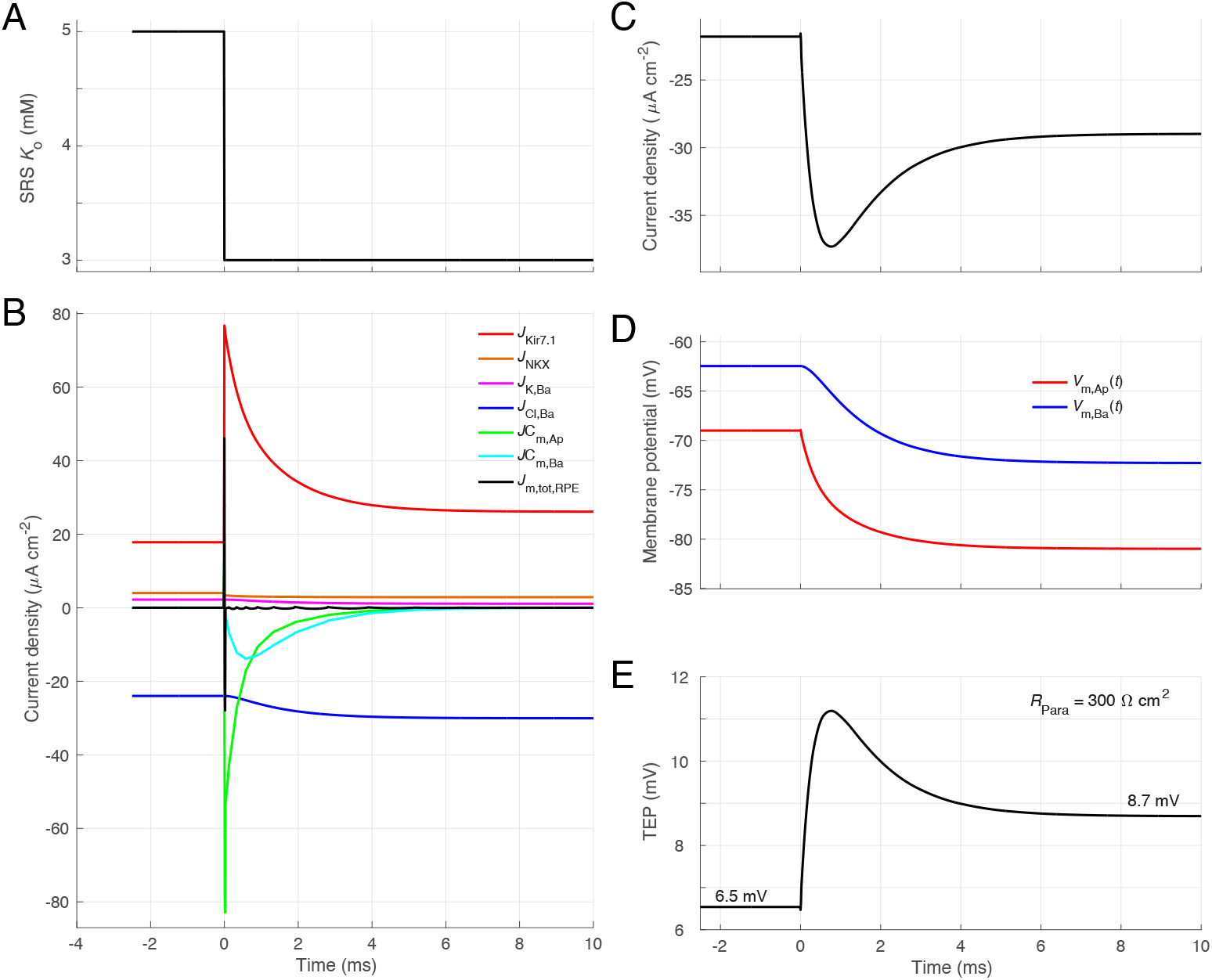
RPE model response to a step decrease in SRS *K*_o_. **A**. Input to model: step decrease in *K*_o_ from 5 mM to 3 mM. **B**. Time course of all membrane currents of the RPE model in response to the step decrease of *K*_o_. The membrane currents (including the apical and basolateral capacitive currents) sum dynamically to zero (black trace), as required for a solution to Eqs 25, 26. The initial conditions of the the differential equations are manifest in the baseline currents, and are the same as those used for the principal model of Fig. 7C (magenta trace), including *J*_NKX, RPE, rest_ = 4 μA cm^-2^, *J*_Kir7.1, rest_ = 17.8 μA cm^-2^, *R*_para_ = 300 Ω cm^-2^ (cf. **Table 2**). RPE cells were assumed to have 3-fold the capacitance density of their geometrical surface to assess whether capacitance would cause a delay in reaching the steady state. **C**. Paracellular current: in addition to Kir7.1 and NKX currents, the paracellular current includes a capacitive contribution. **E**. Response of the apical and basal membrane potentials to the step decrease in SRS *K*_o_. **F**. Transepithelial potential causes by the current flowing through the paracellular resistance; at the steady-state, which is reached in ∼ 7 ms, the steady-state TEP response to the step change in *K*_o_ is 2.2 mV.

**Figure S4.1.**
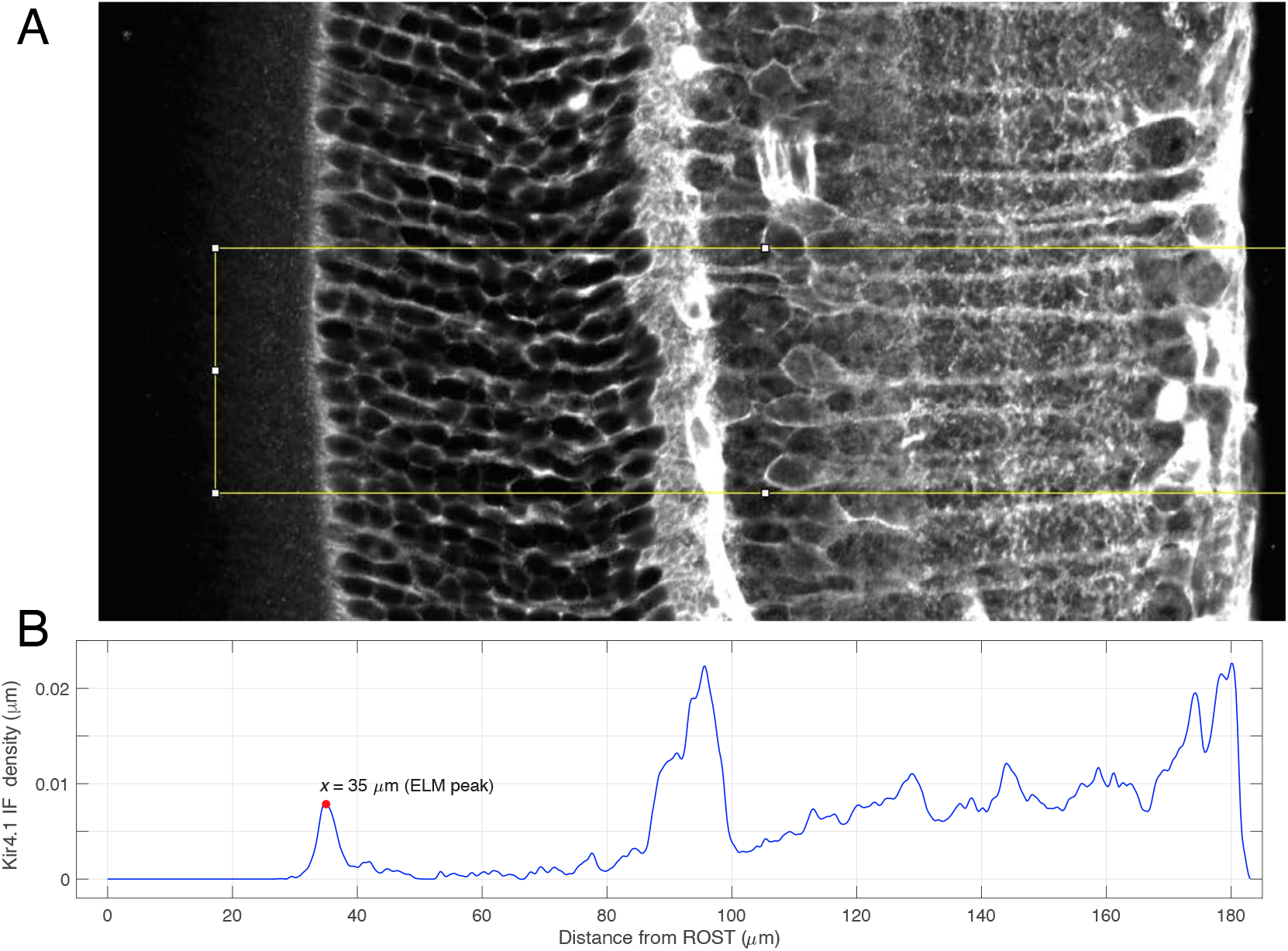
**A**. Cryosection of C57Bl/6 mouse retina stained with a Kir4.1 antibody (anti-rabbit, Alomone Labs, #APC-035, conjugated to Alexa488 dye), imaged with a Nikon A1 confocal (40X). **B**. ImageJ profile plot of the fluorescence in the yellow ROI in A scaled to have unit area, i.e., to create a density function of Kir4.1 expression over the axial coordinate. Faint but reliably detectable immuno-fluorescence is present in the SRS to the left of the ELM peak at 35 μm.

**Figure S4.2.**
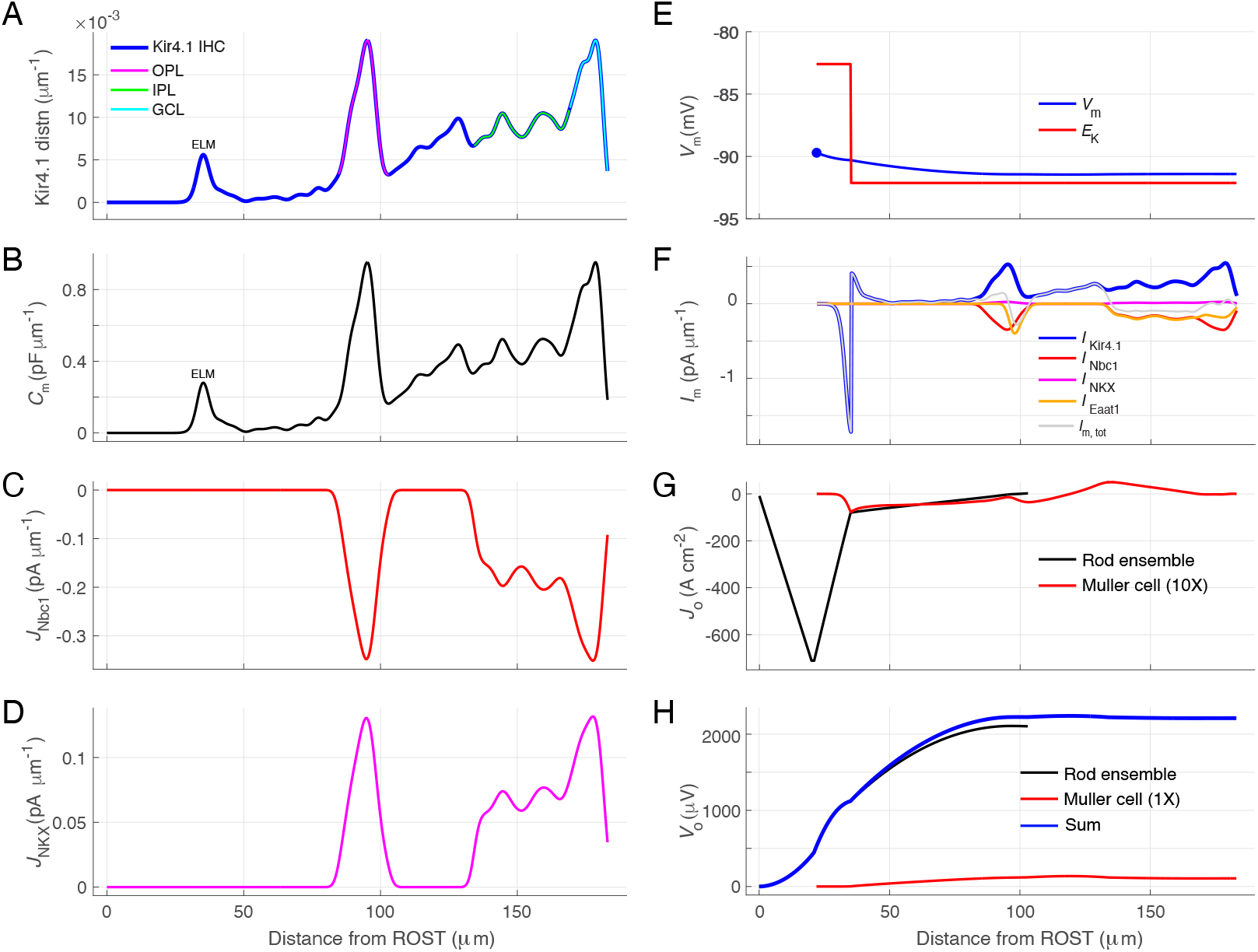
Model derivation of the dark current and field potential of mouse Müller glia cells. **A**. Axial distribution of Kir4.1 immunofluorescence (blue; “IF”), with subregions of the retina identified with overlaid colors on the IF distribution (sampling grid, 0.5 μm). **B**. Axial capacitance distribution of an average Müller glial cell: the distribution in A scaled so that its integral is 50 pF, the measured total capacitance of the mouse Müller cell (Chao et al., 1997). **C**. Axial distribution of Nbc1 (Na^+^/HCO_3_^−^ cotransporter) current; the integral of the current distribution, −14.2 pA, corresponds to the extrusion (as HCO_3_^−^) of all the CO_2_ generated by aerobic respiration needed to sustain the dark current of 22 rods. **D**. Axial distribution of the NKX exchange current (1 pA total). **E**. Axial profiles of the Müller cell membrane potential *V*_m_ and the K^+^ Nernst potential *E*_K_. The step decrease in *E*_K_ at *x* = 35 μm arises from the assumed distribution of *K*_o_ (5 mM in the SRS and 3.5 mM elsewhere (**Fig. 2**). *V*_m_(*x*) is an axial profile that enables the resting model Müller cell to satisfy the Equation of Continuity, i.e., inward and outward current over the cells integrates to zero. **F**. Current density profiles of the three molecular mechanisms, Kir4.1 channels, Nbc1 and NKX. **G**. Extracellular current densities (μA cm^-2^) of rods (black curve) and Müller cells (red curve, displayed at 10× scale). The extracellular limb of the circulating currents of both cell types flow toward the RPE, but that of the rods is much larger, mainly due to the 22:1 ratio of rods to Müller cells. **H**. Field potentials of the rods (black curve) and Müller cells (red) and their sum (blue).

**Figure S4.3.**
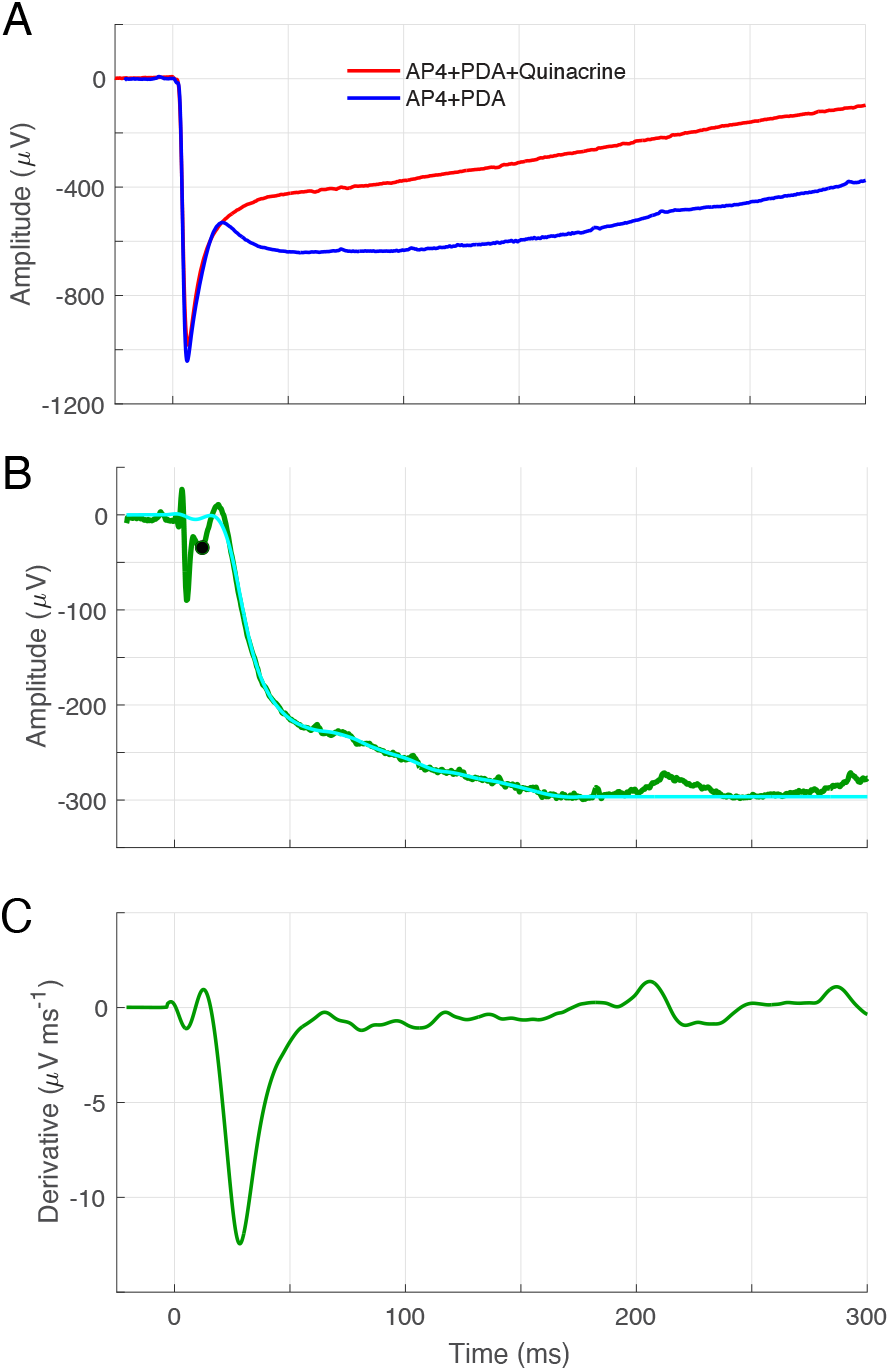
Extraction of the “Slow PIII” component of the ERG from recordings in the presence of AP4 and PDA, with and without quinacrine. **A**. Average ERGs of *Gnat2*^−/–^ mice to a flash delivering ∼ 5×10^5^ photons μm^-2^ **B**. The difference of the traces in A (dark green), and a smoothed version (cyan). On the assumption that quinacrine blocks Kir4.1 channels, the derived “Slow PIII” trace is the trans-retinal field potential generated by the Müller cells. **C**. Derivative of the trace in B (computed with the Matlab “gradient” operator). This serves to emphasize the time window over which the most rapid changes in Kir4.1 current occur. The smoothed difference trace in B (cyan), scaled as indicated, was used as the Müller cell source in Fig. 7B in predicting the ERG from 0 to 3 s for a stimulus producing 3.5×10^5^ photoisomerizations per rod. Thus, the present model of the synaptically blocked ERG *a*- and *c*-waves is semiempirical.

## Notes

### Competing Interest Statement

The authors have declared no competing interest.

